# Loss of Caspase-8 function in combination with SMAC mimetic treatment sensitizes Head and Neck Squamous Carcinoma to radiation through induction of necroptosis

**DOI:** 10.1101/2020.04.17.039040

**Authors:** Burak Uzunparmak, Meng Gao, Antje Lindemann, Kelly Erikson, Li Wang, Eric Lin, Steven J. Frank, Frederico O. Gleber-Netto, Mei Zhao, Heath D. Skinner, Jared Newton, Andrew G. Sikora, Jeffrey N. Myers, Curtis R. Pickering

## Abstract

Caspase-8 (*CASP8*) is one of the most frequently mutated genes in head and neck squamous carcinomas (HNSCC), and mutations of *CASP8* are associated with poor overall survival. The distribution of these mutations in HNSCC suggests that they are likely to be inactivating. Inhibition of *CASP8* has been reported to sensitize cancer cells to necroptosis, a unique cell death mechanism. Here, we evaluated how *CASP8* regulates necroptosis in HSNCC using cell line models and syngeneic mouse xenografts. *In vitro*, knockdown of *CASP8* rendered HNSCCs susceptible to necroptosis induced by a second mitochondria-derived activator of caspase (SMAC) mimetic, Birinapant, when combined with pan-caspase inhibitors Z-VAD-FMK or Emricasan. Strikingly, inhibition of *CASP8* function via knockdown or Emricasan treatment was associated with enhanced radiation killing by Birinapant through induction of necroptosis. In a syngeneic mouse model of oral cancer, Birinapant, particularly when combined with radiation delayed tumor growth and enhanced survival under *CASP8* loss. Exploration of molecular underpinnings of necroptosis sensitivity confirmed that the level of functional receptor-interacting serine/threonine-protein kinase-3 (RIP3), a key enzyme in the necroptosis pathway was crucial in determining susceptibility to this mode of death. Although an *in vitro* screen revealed that many HNSCC cell lines were resistant to necroptosis due to low levels of RIP3, patient tumors maintain RIP3 expression and should therefore remain sensitive. Collectively, these results suggest that targeting the necroptosis pathway with SMAC mimetics, especially in combination with radiation, may be a relevant therapeutic approach in HNSCC with compromised *CASP8* status, provided that RIP3 function is maintained.

**Significance:** *CASP8* status regulates necroptotic death in HNSCC and this pathway can be exploited therapeutically.

## Introduction

Head and Neck Squamous Carcinoma (HNSCC) that comprises epithelial tumors originating from the mucosa of oral cavity, oropharynx, larynx and hypopharynx is one of the most common cancers in the world, with the diagnosis of nearly 650,000 new cases and more than 300,000 cancer related deaths annually (1). 5-year survival rate for HNSCC remains at ∼50% due to resistance to standard-of-care therapy that involves surgery, radiation and/or platinum- or taxane-based chemotherapy, or combination of these modalities (2). Integrative genomic analysis of HNSCC has uncovered that Caspase-8 (*CASP8*) is one of the most frequently mutated genes in HNSCC, with somatic mutations detected in ∼10% of cases (3,4). The distribution of *CASP8* mutations observed in patient tumors and cell lines suggests that they are likely to be inactivating-type mutations where protein function is compromised (4).

*CASP8* is an aspartate-specific cysteine protease that plays a key role in the initiation of extrinsic apoptosis (5). Binding of a death ligand (i.e. TNF-Related Apoptosis-Inducing Ligand [TRAIL]) to its cognate receptor (i.e. TRAIL-receptor) leads to formation of a Death-Inducing Signaling Complex (DISC) at the cytoplasmic tail of the death receptor that comprises the adaptor protein FADD (Fas-Associated with Death Domain) and Procaspase-8. Processing of Procaspase-8 within the DISC yields active *CASP8*, which translocates to the cytosol to cleave and activate its downstream executioner caspases such as Caspases -3 and -7, executing the apoptosis pathway (6–8). Due to the key role it plays in death-receptor mediated apoptosis, *CASP8* has long been considered a tumor suppressor gene (9). This is consistent with the observation that *CASP8* activity is impaired in a variety of cancer types such as neuroblastoma, medullablastoma, and HNSCC through mutations and epigenetic silencing (4,10,11). However, the presence of functional *CASP8* is also crucial for the maintenance of life since *CASP8*^*-/-*^ mice die intranatally around embryonic day 11, resulting from uncontrolled necroptosis (12).

Necroptosis is a unique mechanism of regulated cell death stimulated upon death receptor signaling (i.e. TNFα signaling) that relies on the activation of mixed lineage kinase domain-like (MLKL), a pseudokinase, by receptor-interacting serine/threonine protein kinases-1 and -3 (RIP1 and RIP3). *CASP8* regulates kinase activity of RIP1 and RIP3 both of which contain *CASP8* cleavage sites (13,14). TNFα binding to its cognate receptor, TNFR1, leads to formation of Complex-I that contains TNFR-associated death domain (TRADD), TNFR-associated factor 2 (TRAF2), inhibitor of apoptosis proteins (IAPs) cIAP1/cIAP2, and RIP1. Ubiquitylation of RIP1 by cIAP1/cIAP2 within Complex-I culminates in the activation of the canonical Nuclear Factor-κB (NF-κB) pathway. When cIAPs are inhibited pharmacologically, such as with the second mitochondria-derived activator of caspase (SMAC) mimetic Birinapant, RIP1 recruits *CASP8* to form cytosolic Complex-IIB to initiate apoptosis (15). In cases where *CASP8* is inhibited by chemicals, such as Z-VAD-FMK, *CASP8* regulation over RIP1/RIP3 kinase activity is abrogated that results in the assembly of Complex-IIC in the cytosol, consisting of RIP1, RIP3 and MLKL (16). MLKL is phosphorylated, trimerized and activated within Complex-IIb upon which it translocates to the plasma membrane to induce membrane permeabilization and subsequent necroptotic cell death (17).

SMAC mimetics are small-molecule inhibitors that promote caspase activation and apoptosis through neutralization of IAPs (18). Preclinical studies have highlighted the therapeutic potential of SMAC mimetics through induction of cancer cell death directly (19) or via synergistic interaction with a variety of cytotoxic therapy approaches, including chemotherapy (20,21), radiotherapy (22,23) or immunotherapy (24). The SMAC mimetic Birinapant was found to enhance cytotoxicity induced by death ligands in a panel of HNSCC cell lines (25). Birinapant also synergizes with radiation to prevent tumor growth in various xenograft models of HNSCC bearing genomic amplifications of FADD and cIAP1 *in vivo* (25). Other SMAC mimetic compounds such as LCL161 and ASTX660 have also been shown to confer *in vivo* radiosensitivity to HNSCC xenografts (26,27). However, how mutations and/or loss of *CASP8* impacts necroptosis in HNSCC and whether modulation of the necroptosis pathway with these small molecule agents might have potential clinical utility in the context of *CASP8* loss have largely been unexplored.

In this study, we found that deletion of *CASP8* rendered HNSCCs susceptible to necroptosis induced by the SMAC mimetic Birinapant. Inhibition of *CASP8* function was also associated with enhanced necroptotic killing by radiation when combined with Birinapant *in vitro* and *in vivo*. We further demonstrated that the level of RIP3 expression determines necroptosis sensitivity in HNSCC. These findings provide preclinical justification for use of the necroptosis pathway as a therapeutic target in HNSCC patients.

## Materials and Methods

### HNSCC Cell Lines

Human derived-HNSCC cell lines were maintained as previously described (28). Murine oral cancer (MOC) 1 cell line was provided by Dr. R. Uppaluri (Washington University School of Medicine) (29). All cell lines were authenticated by short tandem repeat (STR) profiling and cultured for no longer than 15 passages before use in experiments.

### Genomic Analysis

The Cancer Genome Atlas (TCGA) data were obtained from the TCGA PanCancerAtlas Project (30) and analyzed for *CASP8* mutations and RIP3 expression for HNSCC. Cell line RIP3 gene expression data is available through the Gene Expression Omnibus (GEO) (accession GSE122512) (31).

### Engineering of stable cell lines

Cell lines were transduced with lentiviral shRNA constructs against *CASP8* (shCASP8) or control scrambled shRNA, containing GFP and puromycin resistance gene (GE Dharmacon). shCASP8 and control cells were GFP-sorted and subjected to puromycin selection (1µg/ml). After antibiotic treatment, control and shCASP8 cells were assessed for protein expression of *CASP8* by Western Blotting (WB). CRISPR/Cas9 sgRNA constructs were designed to knock out *CASP8* in MOC1 cells (32). Knockout of *CASP8* was validated via WB screening and sequencing.

RIP3 was knocked down using lentiviral shRNA constructs (GE Dharmacon) in select MOC1 clonal cell lines carrying WT RIP3, as described above for *CASP8*. Select MOC1 clones that show lack of protein expression of RIP3 were transduced with inducible lentiviral vectors encoding WT (Plasmid #73701) or kinase-dead mutant (D143N) RIP3 (Plasmid #73703) purchased from Addgene (These plasmids were gifted by Dr. Francis Chen). Control pTRIPZ empty vector was obtained from MD Anderson Functional Genomics Core Facility. MOC1 cells transduced with control or RIP3 expression constructs were subjected to puromycin selection (1µg/ml). Expression of WT or D143N RIP3 was induced by Doxycycline (1µg/ml) for 72 hours and verified by WB.

For the *in vivo* experiments, inducible lentiviral shRNA constructs were designed against Luciferase (shLUC) or *CASP8 (*shCASP8) in collaboration with the MD Anderson Institute for Applied Cancer Science as described previously (33). MOC1 cells transduced with shLUC or shCASP8 vectors were subjected to puromycin selection (1µg/ml), treated with Doxycycline (50ng/ml) for 72 hours and verified by WB. shRNA/sgRNA oligo sequences are available in **Supplementary Table S1**.

### Cell proliferation and viability assays

To evaluate cell proliferation, HNSCC cells were plated in 96-well plates at 2 densities: 50cells/well and 100cells/well. Cell density was measured using CellTiter-Glo (Promega). Luminescence reads were taken at the indicated time points and normalized to Day 0 reads to calculate cell doublings.

To assess cell viability following drug treatments, 3-10 × 10^3^ cells were plated in 96-well plates and allowed to attach overnight. Cells were then treated with 0.01% DMSO (mock treatment) or Birinapant, Z-VAD-FMK (or Emricasan), TNFα (or TRAIL) and Necrostatin-1s either alone or in combinations at the indicated doses for 24 hours, after which cell viability was measured by CellTiter-Glo. Average luminescence values taken for each treatment condition were normalized to that of mock-treated cells from the same experiment to calculate % cell density. All treatments were carried out in triplicates or greater. Please refer to **Supplementary Table S2** for detailed information about the drugs/reagents used in the study.

### Clonogenic survival assays

HNSCC cells were seeded in 6-well plates at predetermined densities and allowed to adhere overnight. The next day, cells were treated with 0.01% DMSO (mock treatment) or Birinapant, Z-VAD-FMK (or Emricasan) and Necrostatin-1s either alone or in combinations at the indicated doses. For radiosensitivity assays, treatments described above were followed by exposure to either 2, 4 or 6 Gray (Gy) radiation. 24 hour after treatments, media containing the drug dilutions were aspirated and replaced with fresh media. Colonies were allowed to form for 5-12 days, after which they were fixed in methanol and stained with crystal violet (2%). Wells containing surviving colonies were scanned and colonies with more than 50 cells were counted with the guidance of ImageJ software (NIH, Bethesda, MD). The number of surviving colonies per well was calculated for each treatment condition and normalized to that of mock-treated cells from the same experiment to calculate % colony counts. For the radiosensitivity assays, average surviving colony counts were normalized to those of mock-treated cells of each radiation dose from the same experiment to calculate surviving fractions. *Log10* of surviving fractions were plotted. All treatments were carried out in triplicates.

### Annexin-V assays

Annexin-V staining was performed in accordance with the manufacturer’s guidelines (BD Biosciences). Briefly, 2.5-7.5 × 10^5^ cells were seeded in 6cm dishes and allowed to adhere overnight. The next day, cells were treated as indicated. All treatments were carried out in triplicates. 24 hour after treatments, media was collected from the dishes and set aside to keep any floating cells. Adherent cells were harvested with trypsin and added to the previously collected media. The cells were then centrifuged, washed once in cold Phosphate Buffered Saline (PBS) and stained with Annexin-V APC (BD Biosciences) and 1µmol/L SytoxBlue (ThermoFisher) in Annexin-V binding buffer. Samples were mixed gently and incubated at room temperature for 30min after which they were subjected to flow cytometry analysis. Flow data were analyzed using FlowJo software.

### Western blotting

2.5-7.5 × 10^5^ HNSCC cells were seeded in 6cm dishes and allowed to adhere overnight. The next day, cells were treated as indicated. For radiosensitization studies, cells were exposed to 6Gy radiation. 24 hour after treatments, whole-cell lysates were obtained and western blot analysis was carried out as previously described (4). Please refer to **Supplementary Table S3** for the list of western blot antibodies.

### *In vivo* xenograft model

All *in vivo* experiments were carried out with approval of the Institutional Animal Care and Use Committee (IACUC) at MD Anderson. 2 × 10^6^ MOC1 cells transduced with an inducible shRNA against *CASP8* were injected into the right flank of WT female C57BL/6 mice obtained from Envigo/Harlan Labs in the presence of Matrigel (Corning). 3 days after inoculation, mice were randomized and placed on control (Global 18% Protein Rodent Diet) or DOX diet (doxycycline hyclate added at 625 mg/kg) obtained from Envigo to induce knockdown of *CASP8 in vivo*. Control and shCASP8 mice were randomized into 4 treatment groups (***n=*7-10** mice/each) 27 days after injection of cells, when the average tumor volume reached ∼150 mm^3^. Treatments included Birinapant (15mg/kg Birinapant, every 3 days for 4 weeks, IP), radiation (5 × 2Gy, Monday to Friday for 1 week) or the combination (A detailed *in vivo* treatment schema is available in **Supplementary Fig. S10**). Tumor measurements were taken three times a week, and mice were euthanized when tumor burden has reached >1.5cm in any dimension. Tumor samples were collected from a subset of control and shCASP8 mice that were not recruited in the drug treatment study. These tumor samples were minced and cultured in medium for 48 hours. Cells shed from the tumors that have attached to culture dishes were collected, lysed and subjected to WB analysis for *CASP8*.

### Statistical Analysis

Kaplan-Meier method was used to calculate overall survival for the patient population. *P* values used to compare gene expression levels for necroptosis markers between different patient cohorts were computed using the Mann-Whitney U test. Student *t* tests were conducted to analyze *in vitro* data. For the *in vivo* studies, a two-way ANOVA test was used to make tumor volume comparisons between animal cohorts. Differences in survival rates between groups were compared using the log-rank (Mantel–Cox) test. All data were presented as mean±SD unless otherwise noted. *P* values < 0.05 were considered statistically significant.

## Results

### *CASP8* mutations are associated with radioresistance and poor survival outcomes in HNSCC

Alterations of *CASP8*, most of which are mutations, are found frequently (11.2%) in HNSCCs (4) and are associated with poor overall survival (**Fig. 1A**). To determine whether poor overall survival outcomes might be linked to radioresistance in *CASP8* mutant HNSCCs, we selected a panel of 46 HNSCC cell lines with known *CASP8* status (34). We examined sensitivity to increasing doses of radiation using a standard clonogenic survival assay (35) and determined the level of radiosensitivity using survival fraction at the 2Gy dose (SF2), since 2Gy is the dose typically used in clinical practice to treat HNSCC patients. Interestingly, we found that *CASP8* mutant cell lines were significantly more radioresistant than those with WT *CASP8* (**Fig. 1B**). Since *CASP8* is known to regulate necroptosis and necroptosis can be induced by radiation in a variety of cancers (36,37), we aimed to understand if necroptosis can be exploited to radiosensitize *CASP8* mutant HNSCCs to radiation. We first explored whether necroptosis-related genes are differentially expressed between *CASP8* mutant and WT tumors in TCGA. The top differentially expressed genes include at least 3 necroptosis-related genes. MLKL (mixed lineage kinase domain-like, p=3.39×10^−10^), FASLG (Fas ligand, p=1.97×10^−9^), and TNFRSF10A (TNF Receptor Superfamily Member 10A, TRAIL receptor, DR4, p=2.63×10^−7^) showed higher expression in *CASP8* mutant HPV (-) oral cancers than their WT counterparts, suggesting that necroptosis might be a potential pathway to target in *CASP8* mutant HNSCCs (**Fig 1C**.).

**Figure 1.**
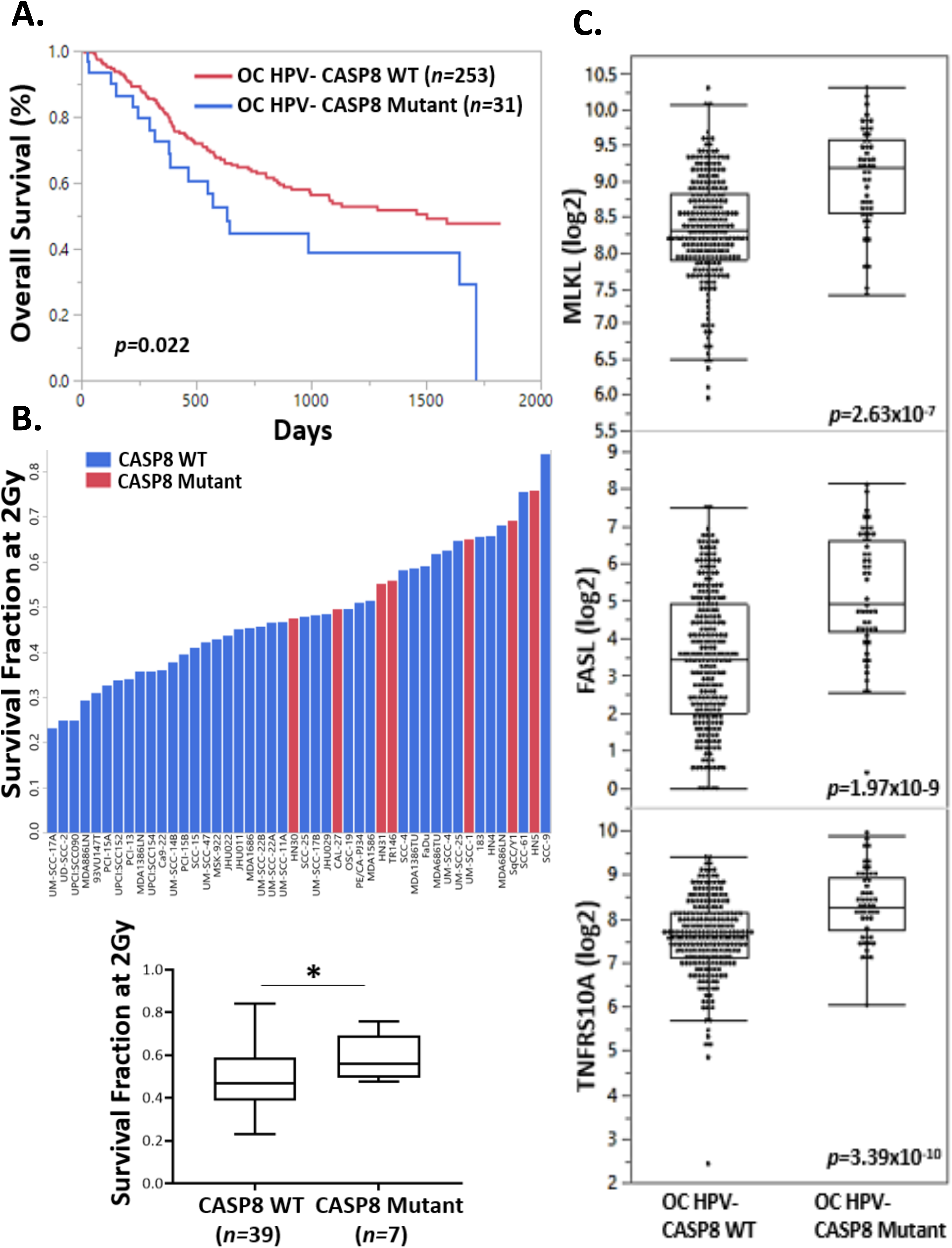
*CASP8* mutations are associated with radioresistance and poor survival outcomes in HNSCCs. **A.** Kaplan-Meier survival plots for *CASP8* in 284 HPV- oral cavity (OC) HNSCCs in TCGA. **B.** A panel of 46 HNSCC cell lines were sequenced for *CASP8* and evaluated for sensitivity to radiation using clonogenic survival assays. Surviving fraction following 2Gy of XRT was used to determine radiosensitivity. (Cell lines with *CASP8* mutation were marked in red). Cell lines were then grouped for *CASP8* status and the average clonogenic survival data were shown for each group. Student t test was used for statistics. *p<0.05 **C**. Scatter plots show gene expression for MLKL, FASL and TNFRSF10A (TRAIL receptor) by *CASP8* status in TCGA HPV- oral cancer samples. Mean values are shown by the bar. P values were computed using Mann-Whitney test.

### Knockdown of *CASP8* sensitizes HNSCCs to necroptotic death by Birinapant and Z-VAD-FMK

Inhibition or mutation of *CASP8* predisposes a wide variety of cancers to necroptosis (38). In an effort to understand how inactivating mutations of *CASP8* that result in loss of protein function impact necroptosis sensitivity in HNSCC, we stably knocked down *CASP8* using a short hairpin RNA (shRNA) in 2 *CASP8* WT HNSCC cell lines, namely the human-derived UMSCC-17A cells and mouse-derived syngeneic oral cancer MOC1 HNSCCs (29). Knockdown of *CASP8* alone did not significantly affect proliferative or colony-forming abilities of the cell lines (**Supplementary Fig. S1**). The UMSCC-17A and MOC1 *CASP8* knockdown and scrambled shRNA control cells (cells transduced with a lentiviral construct lacking the shRNA) were treated with the SMAC mimetic Birinapant alone or in combination with the pan-caspase inhibitor Z-VAD-FMK at concentrations previously shown to be active (25). This combination is a standard method to experimentally induce necroptotic death (39). Cell survival was assessed by cell viability and clonogenic assays. Knockdown of *CASP8* significantly increased the sensitivity of UMSCC-17A and MOC1 cells to single agent Birinapant and Birinapant in combination with Z-VAD-FMK (**Fig. 2A, B and Supplementary Fig. S2-S4**). To confirm that this was a necroptotic death the RIP1 inhibitor, Necrostatin-1s was used. Importantly, reduction in cell viability and clonogenicity was abrogated by Necrostatin- 1s, indicating that the observed mode of cell death was necroptosis not apoptosis (**Fig. 2A, B and Supplementary Fig. S2-S4**). Further indication that this was not an apoptotic death is the inclusion of Z-VAD-FMK treatment, a pan-caspase inhibitor that blocks apoptosis, which enhanced cell death rather than prevented it, consistent with necroptosis.

**Figure 2.**
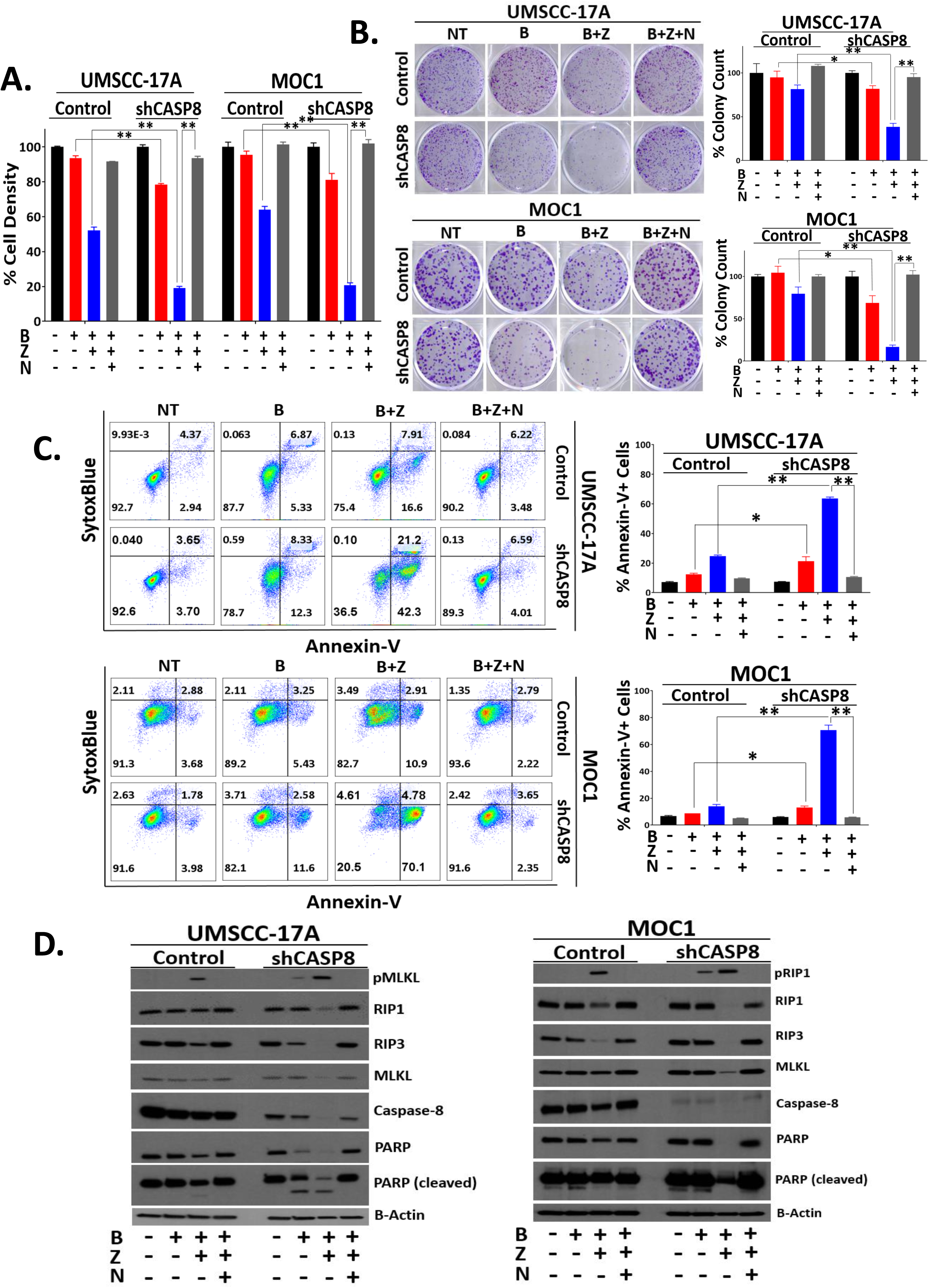
Knockdown of *CASP8* sensitizes HNSCCs to necroptotic death by Birinapant and Z-VAD-FMK. UMSCC-17A and MOC1 control and shRNA knockdown (shCASP8) cells were treated with Birinapant (**B** [200nmol/L for the UMSCC-17A cells; 1μmol/L for the MOC1 cells]), Z- VAD-FMK (**Z** [5μmol/L for both the cell lines]), Necrostatin-1s (**N** [10μmol/L for both the cell lines]) or the combinations as indicated for 24 hours. **A**. Cell viability was assessed using CellTiter-Glo. Values normalized to nontreated cells from the same experiment to calculate % cell density. All treatments were carried out in triplicates. More detailed analysis can be found in **Supplementary Fig. S2A-B** and **Supplementary Fig. S3. B**. Representative images of clonogenic survival assays. 24 hour after treatments (Birinapant doses reduced to 50nmol/L and 250nmol/L for the UMSCC-17A and MOC1 cells respectively, for this assay. Other drugs used at the same concentrations as stated above), drug dilutions were washed out, colonies were allowed to form for 5-12 days, after which they were stained and counted. Surviving colonies were normalized to nontreated cells from the same experiment to calculate % colony count. All treatments were carried out in triplicates. Please refer to **Supplementary Fig. S4** for more detailed version. **C**. AnnexinV-APC/SytoxBlue staining was performed 24 hour after treatments. % Annexin-V positivity was used as a measure to assess cell death. All treatments were carried out in triplicates. More detailed analysis can be found in **Supplementary Fig. S5. D**. Whole cell lysates were collected 24 hour after treatments and subjected to Western blot analysis for the indicated key cell death markers. β-Actin was used as loading control. Student *t* test was used for statistics. **\***, P<0.05; **\*\***, P<0.001 for the indicated pairwise comparisons.

Although studies by other groups indicate that the cytotoxic effects of SMAC mimetics are potentiated by death ligands such as TNFα and TRAIL in HNSCC (25,27), exogenous supply of death ligands was not associated with a significant increase in Birinapant-induced necroptosis in our study (**Supplementary Fig. S2, S3**), a phenomenon that might be explained by the sufficiency of autocrine/paracrine death receptor signaling in the preclinical cell line models that we utilized (40).

Enhanced necroptotic cell death observed under knockdown of *CASP8* was further evaluated with Annexin-V staining (41). Knockdown of *CASP8* led to a significant increase in Annexin-V positive staining following treatment of UMSCC-17A and MOC1 cells with Birinapant alone and Birinapant in combination with Z-VAD-FMK. Necrostatin- 1s reversed positive Annexin-V staining induced by Birinapant in combination with Z-VAD-FMK in both the cell lines, suggesting the predominant occurrence of RIP1 mediated, necroptotic cell death. (**Fig. 2C and Supplementary Fig. S5**).

The expression of key cell death pathway markers following Birinapant alone and Z-VAD-FMK combination treatments in the absence or presence of Necrostatin-1s was evaluated by Western Blotting (**Fig. 2D**). Knockdown of *CASP8* was associated with a significant increase in the protein levels of phospho-MLKL and phopsho-RIP1 following Birinapant alone and Z-VAD-FMK combination treatments in the UMSCC-17A and MOC1 cells, respectively. The increased phosphorylation was accompanied by a significant reduction in the total protein levels of RIP1, RIP3 and MLKL due to their proteosomal degradation at this 24 hour time point (42). Necrostatin-1s reversed the effects of Birinapant alone and the Birinapant and Z-VAD-FMK combination. Taken together, these data clearly demonstrate that loss of *CASP8* sensitizes HNSCCs to Birinapant and Birinapant plus Z-VAD-FMK induced necroptotic cell death *in vitro*.

### Knockdown of *CASP8* enhances the radiosensitizing effects of Birinapant and Z-VAD-FMK through induction of necroptosis

The combination of SMAC mimetic Birinapant and radiation has been shown to be active in HNSCCs with alterations in cell death pathways (25). Given *CASP8* mutant HNSCC cell lines are more radioresistant than their WT counterparts and *CASP8* mutant HNSCCs show alterations in the necroptosis pathway, we sought to understand whether necroptosis could be exploited therapeutically to treat HNSCCs with compromised *CASP8* status. To test this, *CASP8* knockdown and scrambled shRNA control MOC1 cells were treated with increasing doses of radiation in combination with Birinapant and/or Z-VAD-FMK in the absence or presence of Necrostatin-1s, and assessed for clonogenic survival. Knockdown of *CASP8* alone did not significantly impact radiosensitivity (**Fig. 3A and Supplementary Fig. S6**). Interestingly, single agent Birinapant and the Birinapant plus Z-VAD-FMK combination rendered control cells more radiosensitive. Additionally, the radiosensitizing effects of Birinapant and Birinapant plus Z-VAD-FMK were significantly enhanced under knockdown of *CASP8*. The addition of Necrostatin-1s returned all colony counts to comparable levels to those treated with radiation alone, indicating that necroptosis is the underlying mechanism through which the cells were sensitized. These results were further confirmed with assays for Annexin-V (**Fig. 3B**) and cell viability (**Fig. 3C and Supplementary Fig**.**S7**) that demonstrated similar necroptotic radiosensitization. To further elaborate on the mechanism(s) that lead to cell death, we assayed the levels of key cell death proteins by western blot following induction of necroptosis and radiation treatment (**Fig. 3D**). Birinapant when combined with radiation led to phosphorylation of RIP1 in *CASP8* knockdown but not scrambled shRNA control MOC1 cells, a phenomenon which was accompanied by reduction in the protein levels of RIP1, RIP3 and MLKL, indicating activation of the necroptosis pathway. This phenotype was further enhanced by the addition of Z-VAD-FMK to the treatment and reversed by Necrostatin-1s treatment, indicating that the radiosensitizing effects of Birinapant and Birinapant plus Z-VAD-FMK manifest themselves through induction of necroptosis.

**Figure 3.**
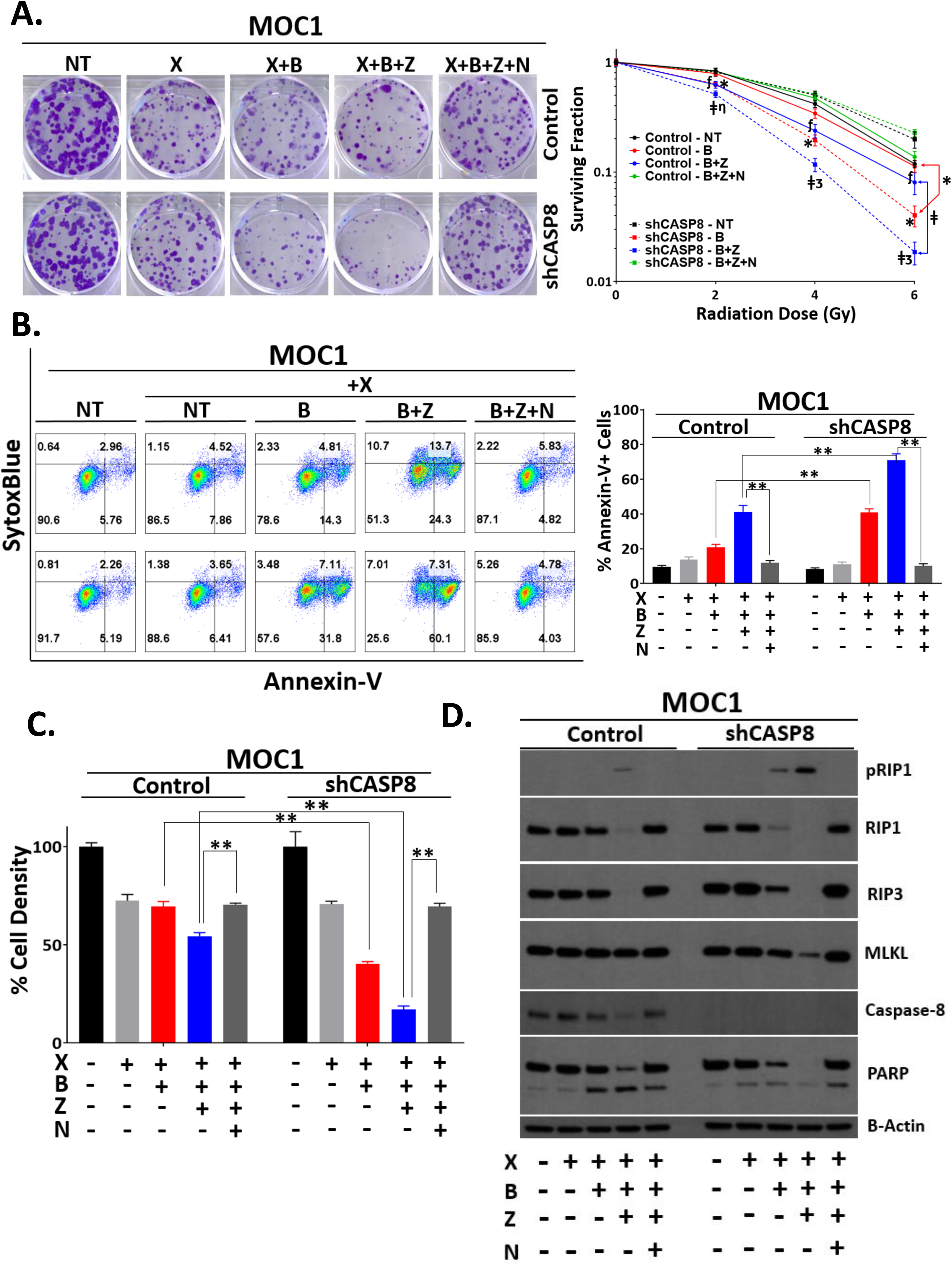
Knockdown of *CASP8* enhances the radiosensitizing effects of Birinapant and Z-VAD-FMK through induction of necroptosis. MOC1 control and shCASP8 cells were treated with radiation (**X** [2, 4 and 6 Gy]), Birinapant (**B** [125nmol/L **in Fig 3A**; 250nmol/L **in Fig3B-3D**]), Z-VAD-FMK (**Z** [5μmol/L]), Necrostatin-1s (**N** [10μmol/L]) or the combinations as indicated. **A**. Representative images of clonogenic survival assays for the **X (4Gy)** conditions. 24 hour after treatments, drug dilutions were washed out, colonies were allowed to form for 5 days, after which they were stained and counted. Surviving colony counts were normalized to nontreated cells (cells treated with no drugs) of each radiation dose from the same experiment. *Log10* of surviving fractions were plotted. All treatments were carried out in triplicates. Student *t* test was used for statistics. **\***, P<0.05; when comparing surviving fractions following X+B treatments between control and shCASP8 cells. **╪**, P<0.05; when comparing surviving fractions following X+B+Z treatments between control and shCASP8 cells. ***f***, P<0.05; when showing reversal of death upon addition of N to X+B+Z for the control cells. **η**, P<0.05; when showing reversal of death upon addition of N to X+B+Z for the shCASP8 cells. **3**, P<0.001; when showing reversal of death upon addition of N to X+B+Z for the shCASP8 cells. Symbols are placed at the radiation doses they refer to. Please refer to **Supplementary Fig. S6** for more detailed version. **B**. AnnexinV-APC/SytoxBlue staining was performed 24 hour after treatments. % Annexin-V positivity was used as a measure to assess cell death. All treatments were carried out in triplicates. **C**. Cell viability was assessed using Cell-Titer Glo. Values normalized to nontreated cells from the same experiment to calculate % cell density. All treatments were carried out in triplicates. More detailed version can be found in **Supplementary Fig. S7**. Student *t* test was used for statistics *, P<0.05; **, P<0.001 for the indicated pairwise comparisons for **Fig. 3B-C. D**. Whole cell lysates were collected 24 hour after treatments and subjected to Western blot analysis for the indicated key cell death markers. β-Actin was used as loading control.

### Susceptibility to necroptosis is determined by levels of RIP3 in HNSCCs

The observation that Z-VAD-FMK enhances Birinapant-induced necroptotic killing under knockdown of *CASP8* might indicate inhibition of residual *CASP8* activity. To further study how loss of *CASP8* impacts necroptosis sensitivity, we genetically deleted *CASP8* in MOC1 cells using CRISPR-Cas9. To rule out clone specific or off-target effects, we designed 2 different single guide RNAs (sgRNA) against *CASP8* to generate multiple independent *CASP8* knockout MOC1 clones. Matching *CASP8* WT clones were created using a non-targeting sgRNA. The MOC1 CRISPR clones were subjected to Western blotting to confirm efficient knockout of *CASP8* along with the parental cell line (**Fig. 4A**). Necroptosis sensitivity was tested in 4 independent *CASP8* WT and *CASP8* knockout clones following treatment with Birinapant or Birinapant in combination with Z-VAD-FMK in the absence or presence of Necrostatin-1s (**Fig. 4B**). Across the clones tested, 2 *CASP8* WT clones (C1 and C2) and 2 *CASP8* knockout clones (g1-1 and g2-1) showed sensitivity to Birinapant plus zVAD-FMK with Necrostatin-1s restoring cell density, indicating a predominantly RIP1-mediated necroptotic cell death. Intriguingly, however, among the *CASP8* knockout clones, clone g2-2 showed complete resistance to Birinapant and Birinapant plus Z-VAD-FMK, and clone g1-4 demonstrated low sensitivity which was not reversed by Necrostatin-1s. These 2 lines exhibited necroptosis resistance despite complete loss of *CASP8*. Additionally, 2 control clones (C4 and C5) also demonstrated resistance to necroptosis, while 1 clone (C1) had enhanced sensitivity to necroptosis. Western blot analysis of the whole-cell lysates obtained from the MOC1 clones and the parental cell line at baseline revealed considerable levels of RIP1 and MLKL for all the cells, but the clones with reduced sensitivity to necroptosis demonstrated lack of protein expression of RIP3 irrespective of *CASP8* status, suggesting that RIP3 levels might determine necroptosis sensitivity in HNSCCs (**Fig. 4C**).

**Figure 4.**
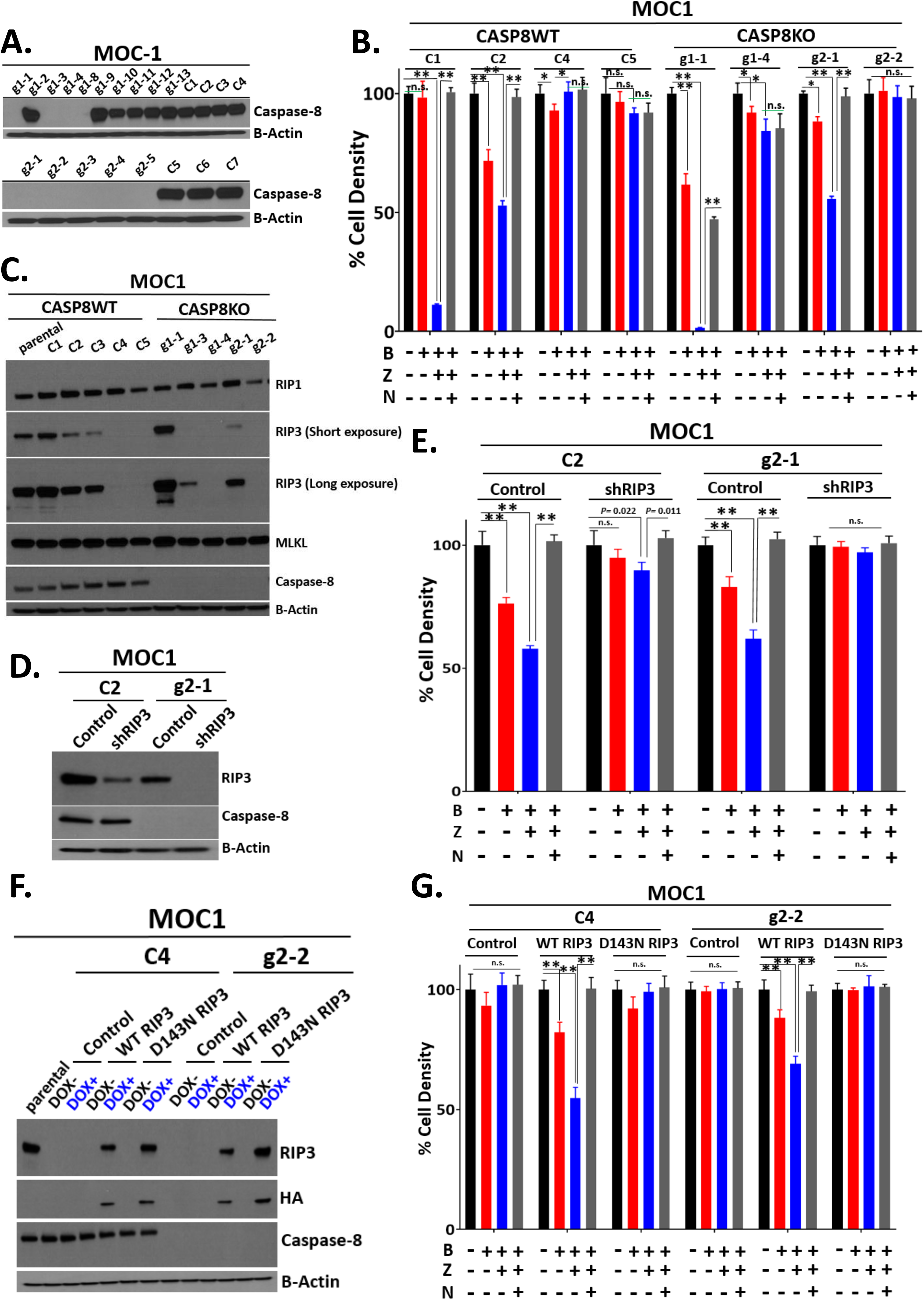
Susceptibility to necroptosis is determined by levels of RIP3 in HNSCCs. **A**. CRISPR/Cas9 was used to knock out *CASP8* in the mouse-derived MOC1 cell line. MOC1 parental cells were transiently transfected with 2 different small guide RNAs designed against mouse *Casp8* (sgRNA-mCASP8 #1 and sgRNA-mCASP8 #2) or a non-targeting sgRNA after which clonal selection/expansion was performed. Engineered clones were subjected to a Western blot screen to identify CASP8WT and CASP8 knockout (CASP8KO) MOC1 clones. **B**. Indicated CASP8WT and CASP8KO MOC1 clones were treated with Birinapant (**B** [1μmol/L]), Z-VAD-FMK (**Z** [5μmol/L]), Necrostatin- 1s (**N** [10μmol/L]) or the combinations for 24 hours. Cell viability was assessed using CellTiter-Glo. Values normalized to nontreated cells from the same experiment to calculate % cell density. All treatments were carried out in triplicates. **C**. Indicated CASP8WT and CASP8KO MOC1 clones were subjected to Western blot analysis for necroptosis markers, RIP1, RIP3 and MLKL. β-Actin was used as loading control. **D**. RIP3 was knocked down using shRNA in 2 Necroptosis sensitive MOC1 clones: the CASP8WT C2 and CASP8KO g2-1 clones. Scrambled shRNA control and shRIP3 cells were subjected to WB to validate knockdown of RIP3. **E**. Control and shRIP3 C2 (CASP8WT) and g2-1 (CASP8KO) MOC1 clones were treated with Birinapant (**B** [1μmol/L]), Z-VAD- FMK (**Z** [5μmol/L]), Necrostatin-1s (**N** [10μmol/L]) or the combinations for 24 hours. Cell viability was assessed by Cell-Titer Glo. **F**. Necroptosis resistant C4 (CASP8WT) and g2- 2 (CASP8KO) MOC1 clones were transduced with control, HA tagged WT RIP3 or HA tagged D143N RIP3 (a kinase domain dead RIP3) inducible expression constructs. RIP3 expression was induced with Doxycycline (50ng/ml). Western blot analysis was performed to validate expression of WT or D143N RIP3 in the indicated MOC1 clones. **G**. Cells engineered in **Fig. 4F** were treated with Birinapant (**B** [1μmol/L]), Z-VAD-FMK (**Z** [5μmol/L]), Necrostatin-1s (**N** [10μmol/L]) or the combinations for 24 hours. Cell viability was assessed by Cell-Titer Glo. Student *t* test was used for statistical analysis. *, P<0.05; **, P<0.001 for the indicated pairwise comparisons.

We next conducted knockdown and overexpression studies to further validate the role of RIP3 in determining necroptosis sensitivity in HNSCCs. shRNA knockdown of RIP3 in 2 necroptosis sensitive MOC1 clones, namely a *CASP8* WT C2 and a *CASP8* knockout g2-1 clone resulted in acquisition of resistance to Birinapant plus Z-VAD-FMK induced necroptotic cell death (**Fig. 4D, 4E**). Conversely, inducible expression of WT but not a kinase dead (D143N) mutant of RIP3 in 2 necroptosis resistant MOC1 clones, namely a *CASP8* WT C4 and a *CASP8* knockout g2-2 clone rendered the cells sensitive to Birinapant plus Z-VAD-FMK induced necroptotic cell death (**Fig. 4F, 4G**). Taken together, these data suggest that the presence of functional RIP3 is necessary for susceptibility to necroptosis in HNSCCs.

### RIP3 expression is silenced in many HNSCC cell lines, but patient tumors show considerable expression

The observation that MOC1 clones lacking functional RIP3 protein show necroptosis resistance led us to test whether RIP3 levels determine necroptosis sensitivity in other HNSCC cell lines. We took a panel of 5 *CASP8* WT and 4 *CASP8* mutant HNSCC cell lines and tested their sensitivity to necroptosis induction and determined the baseline expression of key necroptosis pathway proteins by Western blotting (**Fig. 5A** and **Supplementary Fig. S8**). Consistent with our previous findings, across all the cell lines tested only 3 cell lines that demonstrated considerable baseline protein levels of RIP3, the *CASP8* WT UMSCC-17A and UMSCC-25 cell lines and the *CASP8* mutant HN30 cell line, showed sensitivity to Birinapant plus Z-VAD-FMK which was reversed by Necrostatin-1s. We next evaluated RIP3 gene expression levels in HNSCC cell lines by using RNA sequencing (**Fig. 5B**). RNA expression of RIP3 correlated with RIP3 protein levels in cell lines, and many cell lines demonstrated low expression of RIP3. This suggests that many HNSCC cell lines may be resistant to necroptosis, due to low levels of RIP3. Given our interest in the use of necroptosis as a therapeutic target in HNSCCs, we assessed RIP3 gene expression in HNSCC tumors using the publicly available The Cancer Genome Atlas (TCGA) HNSCC dataset (**Fig. 5C**). Analysis of TCGA tumors confirmed high levels of RIP3 in most tumors, providing justification for therapeutic use of necroptosis in HNSCC. We hypothesize that the loss of RIP3 expression in many cell lines may be due to promoter DNA methylation that occurs *in vitro* (43,44).

**Figure 5.**
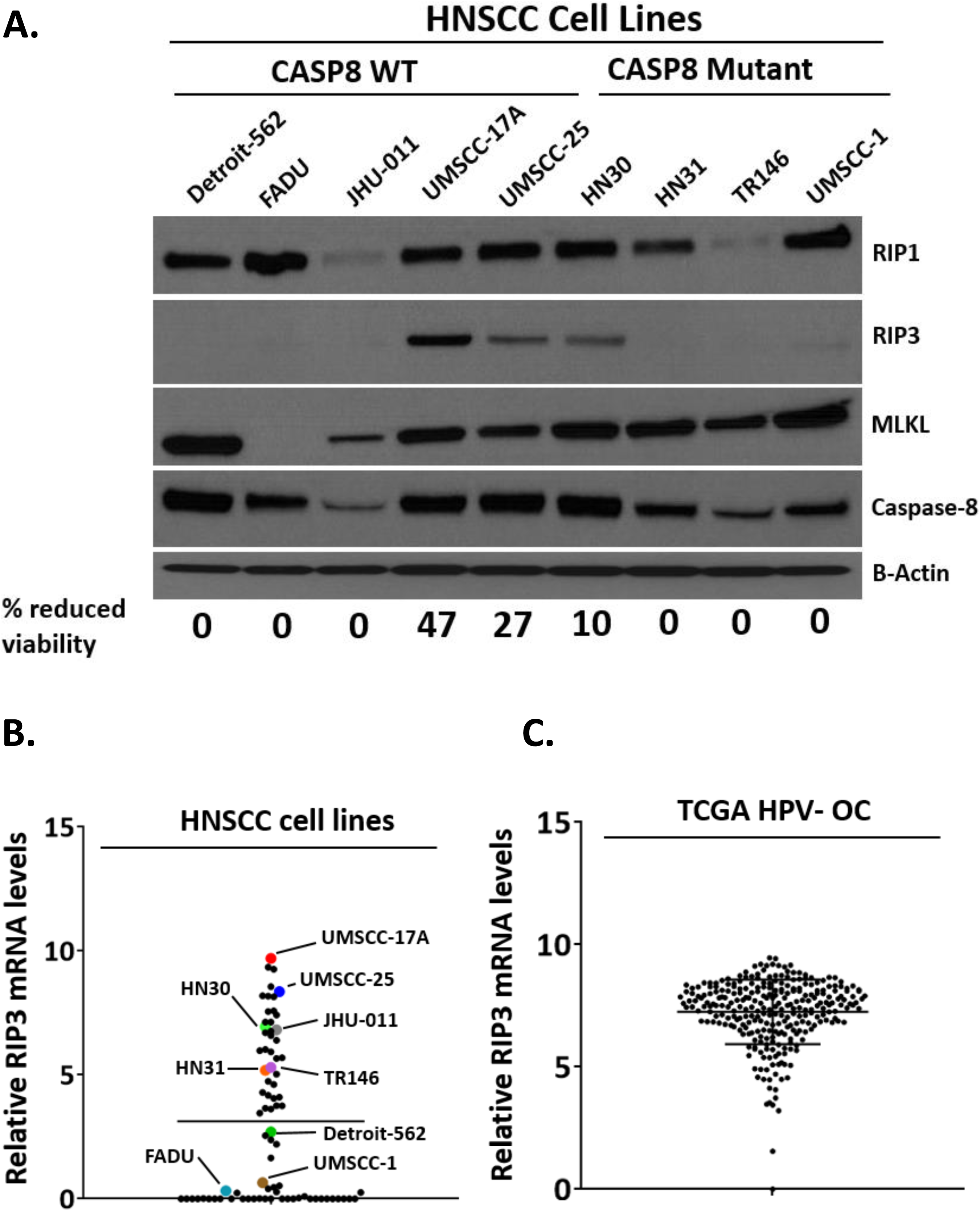
RIP3 expression is silenced in many HNSCC cell lines, but patient tumors show considerable expression. **A**. Cell lysates obtained from a panel of 5 CASP8WT and 4 CASP8mutant human-derived HNSCC cell lines were subjected to Western blot analysis for Caspase-8 and key necroptosis markers. β-Actin was used as loading control. Cell lines were treated with Birinapant (**B** [1μmol/L]), Z-VAD-FMK (**Z** [5μmol/L]), Necrostatin-1s (**N** [10μmol/L]) or the combinations for 24 hours. Cell viability was assessed using Cell-Titer Glo. **“% reduced viability”** was used to assess necroptosis sensitivity for each cell line and calculated based on % reduction in cell density with Birinapant plus Z-VAD-FMK treatment relative to the untreated control that was reversible by Necrostatin-1s. Please refer to **Supplementary Fig. S8** for detailed analysis. **B**. Scatter plot shows gene expression for RIP3 in a panel of 80 human-derived HNSCC cell lines. Mean values are shown by the bar. Cell lines used for the Western Blot analysis were highlighted. **C**. RIP3 gene expression in TCGA HPV- oral cancer samples. Mean values are shown by the bar.

### Loss of *CASP8* increases sensitivity to single agent Birinapant and Birinapant plus radiation *in vivo*

On the basis of our *in vitro* observations that loss of *CASP8* increases Birinapant sensitivity and enhances the radiosensitizing effects of Birinapant in MOC1 cells, we sought to assess the therapeutic efficacy of Birinapant alone or in combination with radiation in the absence or presence of *CASP8* knockdown using the syngeneic MOC1 model *in vivo*. To test this, we generated a MOC1 cell subline transduced with a doxycycline inducible lentiviral *CASP8* shRNA vector (33). Treatment of the *CASP8* shRNA-MOC1 cells with doxycycline, but not vehicle control, led to efficient knockdown of *CASP8 in vitro* (**Supplementary Fig. S9A**). The *CASP8* shRNA-MOC1 cells were injected subcutaneously into the upper leg of syngeneic C57BL/6 mice and knockdown of *CASP8* was achieved *in vivo* by feeding the animals doxycycline containing food or matching control diet throughout the study (**Supplemental Fig. S9B**). Mouse cohorts with control and *CASP8* knockdown MOC1 flank tumors were treated with Birinapant (15mg/kg intraperitoneally) every 3 days for 4 weeks, radiation 2 Gy daily Monday to Friday for 1 week, or the combination when the tumors reached 150 mm^3^ (25). The schema of *in vivo* treatments is shown in **Supplementary Fig. S10**. Treatment with radiation alone significantly inhibited *in vivo* tumor growth (**Fig. 6A, Supplementary Table S4**) and improved survival (**Fig. 6B, Supplementary Table S5**) in both the control and *CASP8* knockdown animal cohorts. However, single agent Birinapant proved effective in delaying tumor growth only when combined with *CASP8* knockdown, a phenomenon which was accompanied by a significant survival benefit. The radiation plus Birinapant combination further reduced *in vivo* tumor growth and provided survival benefit in both the control and *CASP8* knockdown animal cohorts. However, mice bearing *CASP8* knockdown MOC1 tumors demonstrated a significant increase in tumor growth delay and a significantly improved survival when compared to those bearing matching control tumors. Taken together, our results suggest that loss of *CASP8* sensitizes HNSCCs to Birinapant and potentiates its radiosensitizing effects *in vivo*.

**Figure 6.**
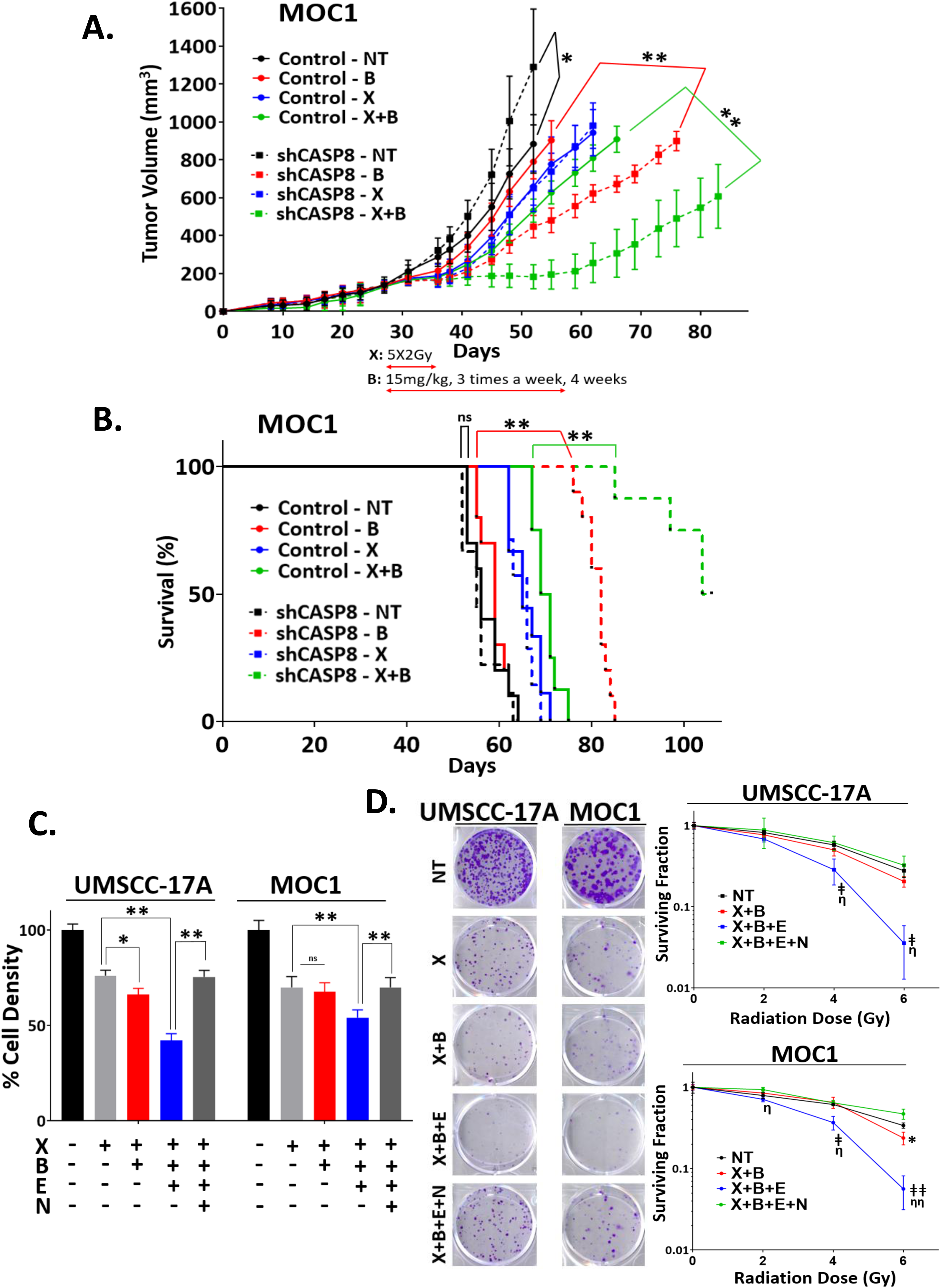
Loss of *CASP8* increases sensitivity to single agent Birinapant and Birinapant plus radiation *in vivo*. **A**. 2 × 106 MOC1 cells transduced with an inducible shRNA against *CASP8* were injected into the right flank of WT female C57BL/6 mice. Mice were randomized and placed on control or DOX diet (doxycycline hyclate added at 625 mg/kg) 3 days post injection to induce knockdown of *CASP8 in vivo* (Please refer to **Supplementary Fig. S9** for the WB images). Control and *CASP8* knockdown mice were randomized into 4 treatment groups (vehicle control, 15mg/kg Birinapant, 5X2Gy radiation or combination, n=7-10/each) 27 days post inoculation when the average tumor volume reached ∼150 mm3. Solid and dashed lines were used to represent control and *CASP8* knockdown mice respectively for the indicated treatment groups. Radiation (**X**) started on Day 27: 2Gy of radiation given Monday to Friday for 1 week (Solid and dashed blue lines). Birinapant (**B**) started on Day 27: 15mg/kg Birinapant given intraperitoneally every 3 days for 4 weeks (Solid and dashed red lines). Black and green lines show vehicle control (**NT**) and combo (**X+B**) groups respectively. A more detailed treatment schema is available in **Supplementary Figure S10**. Error bars represent standard deviation. 2-way ANOVA was used for statistical analysis. **\***p < 0.05, **\*\***p < 0.001 for the indicated pairwise comparisons. **B**. Kaplan-Meier survival curves representing each treatment group. Log-rank (Mantel-Cox) test was used for statistical analysis. **\***p < 0.05, **\*\***p < 0.001 for the indicated pairwise comparisons. **C**. UMSCC-17A and MOC1 parental cells were treated with radiation (**X** [2, 4 and 6 Gy]), Birinapant (**B** [50nmol/L for the UMSCC-17A cells; 250nmol/L for the MOC1 cells]), Emricasan (E [1μmol/L for both the cell lines]), Necrostatin-1s (**N** [10μmol/L for both the cell lines]) or the combinations as indicated for 24 hours. Cell viability was assessed using CellTiter-Glo. Values normalized to nontreated cells from the same experiment to calculate % cell density. All treatments were carried out in triplicates. Student t test was used for statistics. *, P<0.05; **, P<0.001 for the indicated pairwise comparisons. More detailed analysis can be found in **Supplementary Fig. S11**. D. Representative images of clonogenic survival assays for the **X (6Gy)** conditions. 24 hour after treatments (Birinapant doses reduced to 25nmol/L and 125nmol/L for the UMSCC- 17A and MOC1 cells respectively, for this assay. Other drugs used at the same concentrations as stated above), drug dilutions were washed out, colonies were allowed to form for 5-12 days, after which they were stained and counted. Surviving colony counts were normalized to nontreated cells (cells treated with no drugs) of each radiation dose from the same experiment. *Log10* of surviving fractions were plotted. All treatments were carried out in triplicates. Student t test was used for statistics. **\***, P<0.05; when comparing X+B to X alone for the indicated radiation dose. **╪**, P<0.05 and **╪╪**, P<0.001; when comparing X+B+E to X alone for the indicated radiation doses. **η**, P<0.05 and **ηη**, P<0.001; when comparing X+B+E to X+B+E+N for the indicated radiation doses. Please refer to **Supplementary Fig. S12** for more detailed version.

### Chemical inhibition of *CASP8* sensitizes to Birinapant plus radiation

Since only a subset of HNSCC contain inactivation of *CASP8* by mutation we sought to determine whether chemical inhibition of *CASP8* could mimic knockdown of *CASP8* for necroptotic radiosensitization. Emricasan (IDN-655) is a potent pan-caspase inhibitor that was well tolerated in patients when it was tested to reduce liver toxicity in patients with liver diseases (45). We found that Emricasan sensitized *CASP8* WT MOC1 and UMSCC-17A cells to necroptosis when combined with XRT and Birinapant (**Fig 6C, 6D and Supplementary Fig**.**S11, S12**), suggesting that targeting the necroptosis pathway through chemical inhibition of *CASP8* could be a therapeutic strategy to radiosensitize HNSCCs.

## Discussion

In this study, we found that *CASP8* status regulates necroptotic death in HNSCC, and SMAC mimetic treatment may be useful to exploit this pathway for therapeutic benefit. SMAC mimetics have shown therapeutic potential in a variety of cancers, including HNSCC, through induction of cancer cell death directly or via synergistic interaction with other cytotoxic therapies such as chemotherapy, radiotherapy or immunotherapies (18–24). Previous studies of HNSCC have reported that SMAC mimetics, including Birinapant, might synergize with radiation to delay tumor growth in various xenograft models (25–27). We show for the first time that inhibition of *CASP8* function can lead to enhanced radiosensitization by Birinapant through induction of necroptotic death (**Fig 3A-D, Supplementary Fig. S6, S7**). Since *CASP8* mutant HNSCCs might be more radioresistant than their WT counterparts (**Fig 1B**), combining SMAC mimetics with radiation is a potentially promising therapeutic strategy to improve radiation response in HNSCCs with compromised *CASP8* status, and this combination should be further investigated in future studies.

While previous reports have demonstrated that SMAC mimetics alone or in combination with radiation can suppress tumor cell growth in *CASP8* WT HNSCC through induction of apoptosis (25,27), we did not identify a large apoptotic component in most of the cell lines we analyzed. Rather, the apoptotic inhibitor Z-VAD-FMK enhanced cell death in our studies. This discrepancy may be due to the use of different HNSCC cell lines that reflect the genomic diversity of HNSCC. However, it may also indicate the broad therapeutic potential of these treatment combinations. It is likely that many HNSCC may be sensitive to some type of cell death induced by a SMAC mimetic alone or in combination with radiation. The *CASP8* status and other genomic alterations (FADD/BIRC2/BIRC3 amplification, RIP3 expression) may determine whether the death is necroptotic or apoptotic, but many genotypes will be sensitive. Additionally, we found that the mode of cell death for *CASP8* WT cells can be pushed toward necroptosis by adding treatment with a caspase inhibitor, such as Emricasan. The ability to tailor the mode of cell death could facilitate therapeutic synergy with other treatment agents, including immunotherapy.

Immunotherapy is an exciting new treatment modality for HNSCC and many other tumor types, however, only a minority of patients respond. Some tumors seems to have an immunologically cold microenvironment that prevents an immunologic response (46). It has been argued, and recently demonstrated (47), that activation of necroptotic death can potentiate antitumor immunity. Necroptosis is a more immunogenic form of cell death, which could be detrimental to healthy tissues but may be useful as a treatment for cancer. Therefore, it is attractive to speculate that the treatments we have investigated could enhance responses to immunotherapy by inducing an immunogenic necroptotic death.

Caspase inhibition has generally not been thought of as a useful therapeutic approach for cancer because the goal is to promote cell death rather than block it. However, a previous report showed that the caspase inhibitor Emricasan renders clinically relevant models of acute myeloid leukemia (AML) susceptible to Birinapant induced necroptosis *in vitro* and *in vivo* (48). In our study, Emricasan significantly enhanced the radiosensitizing effects of Birinapant in two preclinical models of HNSCC, through induction of necroptosis (**Fig. 6C-D, Supplementary Fig. S11, S12**). Emricasan is well tolerated in patients and is being tested in humans for the treatment of liver diseases characterized by hepatic inflammation and fibrosis (49). It is interesting to propose the combination of Birinapant (or other SMAC mimetics such as ASTX660 or LCL161) and radiation with Emricasan to inhibit apoptotic death but promote necroptotic death, thereby using caspase inhibition for cancer therapy.

Presence of RIP3 has been shown to be pivotal in determining the sensitivity of a variety of cancer types to necroptosis (39). In that study, the cancer cell lines that showed lack of RIP3 expression were found to be resistant to the combination of TNFα, Z-VAD-FMK and the SMAC mimetic SM-164. Similarly, in another study where 8 colon cancer cell lines were tested for sensitivity to a TNFα, SMAC mimetic and Z-VAD-FMK combination, only those that were devoid of RIP3 at the mRNA and protein levels failed to undergo necroptosis (50). Consistent with these results, our data suggest that RIP3 loss can be an underlying mechanism by which HNSCCs become resistant to necroptotic death stimulated with Birinapant and Z-VAD-FMK (**Fig 4A-G**). Loss of protein expression of RIP3 in necroptosis-resistant HNSCCs coincided with low mRNA levels of *Rip3* (**Fig 5A-B**), indicating a likely transcriptional regulation of RIP3. Intriguingly, in a study where mechanisms of RIP3 loss were investigated in various cancer cell lines, treatment of RIP3 lacking cells with the hypomethylating agent 5-aza-2′-deoxycytidine but not the proteosome inhibitor MG132 restored RIP3 expression. Further analyses conducted by the authors revealed that RIP3 loss in those cells was associated with methylation of 4 CpG islands located downstream of the transcription start site (TSS) of *Rip3* (43). Therefore, it is likely that DNA methylation is the underlying mechanism for loss of RIP3 in the preclinical models of HNSCC that we employed in our studies. Since cell lines grown in 2D culture are prone to increased DNA methylation (44), we evaluated RIP3 gene expression in HNSCC tumors using TCGA HNSCC dataset (**Fig. 5C**). Analysis of TCGA tumors revealed that patient tumors show high levels of *Rip3*, providing justification for exploitation of necroptosis therapeutically in HNSCC, and suggesting that the silencing of RIP3 in some cell lines may be an artifact of 2D culture.

In conclusion, here we demonstrate that inhibition of *CASP8* function enhances sensitivity of HNSCCs to Birinapant and Birinapant plus radiation through induction of necroptosis *in vitro* and *in vivo*, on the condition that RIP3 function is maintained. These results provide a strong clinical relevance for the combination of SMAC mimetics like Birinapant and radiation in *CASP8* mutant HNSCCs, a therapeutic approach that might potentially be effective even in *CASP8* wild-type patients with the use of a clinically tolerable caspase inhibitor, such as Emricasan. Further studies to identify optimal and effective combination dose of Emricasan with Birinapant and/or radiation *in vivo* are warranted as are combinations with immunotherapy.

## Abbreviations

HNSCC: head and neck squamous cell carcinoma
OSCC: oral squamous cell carcinoma
*CASP8*: Caspase-8
MOC1: mouse oral cancer-1
cIAP1/2: cellular inhibitor of apoptosis proteins-1/2
SMAC: second mitochondria-derived activator of caspase
RIP1: receptor-interacting serine/threonine-protein kinase-1
RIP3: receptor-interacting serine/threonine-protein kinase-3
MLKL: mixed lineage kinase domain-like
Z-VAD-FMK: carbobenzoxy-valyl-alanyl-aspartyl-[O-methyl]- fluoromethylketone
XRT: radiation therapy

## References

1. Bray F, Ferlay J, Soerjomataram I, Siegel RL, Torre LA, Jemal A. Global cancer statistics 2018: GLOBOCAN estimates of incidence and mortality worldwide for 36 cancers in 185 countries. CA Cancer J Clin 2018;68:394–424.

2. Ang KK, Harris J, Wheeler R, Weber R, Rosenthal DI, Nguyen-Tân PF, et al. Human papillomavirus and survival of patients with oropharyngeal cancer. N Engl J Med 2010;363:24–35.

3. Agrawal N, Frederick MJ, Pickering CR, Bettegowda C, Chang K, Li RJ, et al. Exome sequencing of head and neck squamous cell carcinoma reveals inactivating mutations in NOTCH1. Science 2011;333:1154–7.

4. Pickering CR, Zhang J, Yoo SY, Bengtsson L, Moorthy S, Neskey DM, et al. Integrative genomic characterization of oral squamous cell carcinoma identifies frequent somatic drivers. Cancer Discov 2013;3:770–81.

5. Muzio M, Chinnaiyan AM, Kischkel FC, O’Rourke K, Shevchenko A, Ni J, et al. FLICE, a novel FADD-homologous ICE/CED-3-like protease, is recruited to the CD95 (Fas/APO-1) death--inducing signaling complex. Cell 1996;85:817–27.

6. Chinnaiyan AM, O’Rourke K, Tewari M, Dixit VM. FADD, a novel death domain-containing protein, interacts with the death domain of Fas and initiates apoptosis. Cell 1995;81:505–12.

7. Fu T-M, Li Y, Lu A, Li Z, Vajjhala PR, Cruz AC, et al. Cryo-EM Structure of Caspase-8 Tandem DED Filament Reveals Assembly and Regulation Mechanisms of the Death-Inducing Signaling Complex. Mol Cell 2016;64:236–50.

8. Keller N, Mares J, Zerbe O, Grütter MG. Structural and biochemical studies on procaspase-8: new insights on initiator caspase activation. Structure 2009;17:438–48.

9. Jäger R, Zwacka RM. The enigmatic roles of caspases in tumor development. Cancers (Basel) 2010;2:1952–79.

10. Teitz T, Wei T, Valentine MB, Vanin EF, Grenet J, Valentine VA, et al. Caspase 8 is deleted or silenced preferentially in childhood neuroblastomas with amplification of MYCN. Nat Med 2000;6:529–35.

11. Pingoud-Meier C, Lang D, Janss AJ, Rorke LB, Phillips PC, Shalaby T, et al. Loss of caspase-8 protein expression correlates with unfavorable survival outcome in childhood medulloblastoma. Clin Cancer Res 2003;9:6401–9.

12. Kaiser WJ, Upton JW, Long AB, Livingston-Rosanoff D, Daley-Bauer LP, Hakem R, et al. RIP3 mediates the embryonic lethality of caspase-8-deficient mice. Nature 2011;471:368–72.

13. Lin Y, Devin A, Rodriguez Y, Liu ZG. Cleavage of the death domain kinase RIP by caspase-8 prompts TNF-induced apoptosis. Genes Dev 1999;13:2514–26.

14. Feng S, Yang Y, Mei Y, Ma L, Zhu D, Hoti N, et al. Cleavage of RIP3 inactivates its caspase-independent apoptosis pathway by removal of kinase domain. Cell Signal 2007;19:2056–67.

15. Darding M, Feltham R, Tenev T, Bianchi K, Benetatos C, Silke J, et al. Molecular determinants of Smac mimetic induced degradation of cIAP1 and cIAP2. Cell Death Differ 2011;18:1376–86.

16. Dillon CP, Weinlich R, Rodriguez DA, Cripps JG, Quarato G, Gurung P, et al. RIPK1 blocks early postnatal lethality mediated by caspase-8 and RIPK3. Cell 2014;157:1189–202.

17. Tummers B, Green DR. Caspase-8: regulating life and death. Immunol Rev 2017;277:76–89.

18. Fulda S. Promises and Challenges of Smac Mimetics as Cancer Therapeutics. Clin Cancer Res 2015;21:5030–6.

19. Petersen SL, Wang L, Yalcin-Chin A, Li L, Peyton M, Minna J, et al. Autocrine TNFalpha signaling renders human cancer cells susceptible to Smac-mimetic-induced apoptosis. Cancer Cell 2007;12:445–56.

20. Probst BL, Liu L, Ramesh V, Li L, Sun H, Minna JD, et al. Smac mimetics increase cancer cell response to chemotherapeutics in a TNF-α-dependent manner. Cell Death Differ 2010;17:1645–54.

21. Dineen SP, Roland CL, Greer R, Carbon JG, Toombs JE, Gupta P, et al. Smac mimetic increases chemotherapy response and improves survival in mice with pancreatic cancer. Cancer Res 2010;70:2852–61.

22. Vellanki SHK, Grabrucker A, Liebau S, Proepper C, Eramo A, Braun V, et al. Small-molecule XIAP inhibitors enhance gamma-irradiation-induced apoptosis in glioblastoma. Neoplasia 2009;11:743–52.

23. Giagkousiklidis S, Vellanki SH, Debatin K-M, Fulda S. Sensitization of pancreatic carcinoma cells for gamma-irradiation-induced apoptosis by XIAP inhibition. Oncogene 2007;26:7006–16.

24. Bake V, Roesler S, Eckhardt I, Belz K, Fulda S. Synergistic interaction of Smac mimetic and IFNα to trigger apoptosis in acute myeloid leukemia cells. Cancer Lett 2014;355:224–31.

25. Eytan DF, Snow GE, Carlson S, Derakhshan A, Saleh A, Schiltz S, et al. SMAC Mimetic Birinapant plus Radiation Eradicates Human Head and Neck Cancers with Genomic Amplifications of Cell Death Genes FADD and BIRC2. Cancer Res 2016;76:5442–54.

26. Yang L, Kumar B, Shen C, Zhao S, Blakaj D, Li T, et al. LCL161, a SMAC-mimetic, Preferentially Radiosensitizes Human Papillomavirus-negative Head and Neck Squamous Cell Carcinoma. Mol Cancer Ther 2019;18:1025–35.

27. Xiao R, An Y, Ye W, Derakhshan A, Cheng H, Yang X, et al. Dual Antagonist of cIAP/XIAP ASTX660 Sensitizes HPV- and HPV+ Head and Neck Cancers to TNFα, TRAIL, and Radiation Therapy. Clin Cancer Res 2019;25:6463–74.

28. Zhao M, Sano D, Pickering CR, Jasser SA, Henderson YC, Clayman GL, et al. Assembly and initial characterization of a panel of 85 genomically validated cell lines from diverse head and neck tumor sites. Clin Cancer Res 2011;17:7248–64.

29. Judd NP, Winkler AE, Murillo-Sauca O, Brotman JJ, Law JH, Lewis JS, et al. ERK1/2 regulation of CD44 modulates oral cancer aggressiveness. Cancer Res 2012;72:365–74.

30. Campbell JD, Yau C, Bowlby R, Liu Y, Brennan K, Fan H, et al. Genomic, Pathway Network, and Immunologic Features Distinguishing Squamous Carcinomas. Cell Rep 2018;23:194-212.e6.

31. Gleber-Netto FO, Rao X, Guo T, Xi Y, Gao M, Shen L, et al. Variations in HPV function are associated with survival in squamous cell carcinoma. JCI insight 2019;4.

32. Ran FA, Hsu PD, Wright J, Agarwala V, Scott DA, Zhang F. Genome engineering using the CRISPR-Cas9 system. Nat Protoc 2013;8:2281–308.

33. Sigl R, Ploner C, Shivalingaiah G, Kofler R, Geley S. Development of a multipurpose GATEWAY-based lentiviral tetracycline-regulated conditional RNAi system (GLTR). PLoS One 2014;9:e97764.

34. Kalu NN, Mazumdar T, Peng S, Shen L, Sambandam V, Rao X, et al. Genomic characterization of human papillomavirus-positive and -negative human squamous cell cancer cell lines. Oncotarget 2017;8:86369–83.

35. Skinner HD, Sandulache VC, Ow TJ, Meyn RE, Yordy JS, Beadle BM, et al. TP53 disruptive mutations lead to head and neck cancer treatment failure through inhibition of radiation-induced senescence. Clin Cancer Res 2012;18:290–300.

36. Nehs MA, Lin C-I, Kozono DE, Whang EE, Cho NL, Zhu K, et al. Necroptosis is a novel mechanism of radiation-induced cell death in anaplastic thyroid and adrenocortical cancers. Surgery 2011;150:1032–9.

37. Wang H-H, Wu Z-Q, Qian D, Zaorsky NG, Qiu M-H, Cheng J-J, et al. Ablative Hypofractionated Radiation Therapy Enhances Non-Small Cell Lung Cancer Cell Killing via Preferential Stimulation of Necroptosis In Vitro and In Vivo. Int J Radiat Oncol Biol Phys 2018;101:49–62.

38. Zhou W, Yuan J. SnapShot: Necroptosis. Cell 2014;158:464-464.e1.

39. Najafov A, Zervantonakis IK, Mookhtiar AK, Greninger P, March RJ, Egan RK, et al. BRAF and AXL oncogenes drive RIPK3 expression loss in cancer. PLoS Biol 2018;16:e2005756.

40. Hannes S, Abhari BA, Fulda S. Smac mimetic triggers necroptosis in pancreatic carcinoma cells when caspase activation is blocked. Cancer Lett 2016;380:31–8.

41. Lee E-W, Kim J-H, Ahn Y-H, Seo J, Ko A, Jeong M, et al. Ubiquitination and degradation of the FADD adaptor protein regulate death receptor-mediated apoptosis and necroptosis. Nat Commun 2012;3:978.

42. Cai Z, Jitkaew S, Zhao J, Chiang H-C, Choksi S, Liu J, et al. Plasma membrane translocation of trimerized MLKL protein is required for TNF-induced necroptosis. Nat Cell Biol 2014;16:55–65.

43. Koo G-B, Morgan MJ, Lee D-G, Kim W-J, Yoon J-H, Koo JS, et al. Methylation-dependent loss of RIP3 expression in cancer represses programmed necrosis in response to chemotherapeutics. Cell Res 2015;25:707–25.

44. Morgan MJ, Kim Y-S. The serine threonine kinase RIP3: lost and found. BMB Rep 2015;48:303–12.

45. Pockros PJ, Schiff ER, Shiffman ML, McHutchison JG, Gish RG, Afdhal NH, et al. Oral IDN-6556, an antiapoptotic caspase inhibitor, may lower aminotransferase activity in patients with chronic hepatitis C. Hepatology 2007;46:324–9.

46. Bonaventura P, Shekarian T, Alcazer V, Valladeau-Guilemond J, Valsesia-Wittmann S, Amigorena S, et al. Cold Tumors: A Therapeutic Challenge for Immunotherapy. Front Immunol 2019;10:168.

47. Snyder AG, Hubbard NW, Messmer MN, Kofman SB, Hagan CE, Orozco SL, et al. Intratumoral activation of the necroptotic pathway components RIPK1 and RIPK3 potentiates antitumor immunity. Sci Immunol 2019;4.

48. Brumatti G, Ma C, Lalaoui N, Nguyen N-Y, Navarro M, Tanzer MC, et al. The caspase-8 inhibitor emricasan combines with the SMAC mimetic birinapant to induce necroptosis and treat acute myeloid leukemia. Sci Transl Med. 2016;8:339ra69.

49. Mehta G, Rousell S, Burgess G, Morris M, Wright G, McPherson S, et al. A Placebo-Controlled, Multicenter, Double-Blind, Phase 2 Randomized Trial of the Pan-Caspase Inhibitor Emricasan in Patients with Acutely Decompensated Cirrhosis. J Clin Exp Hepatol 2018;8:224–34.

50. Yang C, Li J, Yu L, Zhang Z, Xu F, Jiang L, et al. Regulation of RIP3 by the transcription factor Sp1 and the epigenetic regulator UHRF1 modulates cancer cell necroptosis. Cell Death Dis 2017;8:e3084.

