## Supplemental figures for "Loss of Caspase-8 function in combination with SMAC mimetic treatment sensitizes Head and Neck Squamous Carcinoma to radiation through induction of necroptosis"

**SUPPLEMENTARY FIGURES (Uzunparmak *et. al.*)**

**Figure S1.** Effects of knockdown of *CASP8* on *in vitro* cell proliferation and clonogenicity in HNSCCs

**Figure S2.** Effects of treatment with Birinapant and Z-VAD-FMK with or without TNFα on cell viability in HNSCCs under loss of *CASP8*

**Figure S3.** Effects of treatment with Birinapant and Z-VAD-FMK with or without TRAIL on cell viability in HNSCCs under loss of *CASP8*

**Figure S4.** Effects of treatment with Birinapant and Z-VAD-FMK on clonogenicity in HNSCCs under loss of *CASP8*

**Figure S5.** Necroptotic effects of treatment with Birinapant and Z-VAD-FMK are enhanced under loss of *CASP8* in HNSCCs.

**Figure S6.** Loss of *CASP8* increases the radiosensitizing effects of Birinapant or Birinapant plus Z-VAD-FMK through induction of necroptosis in HNSCC.

**Figure S7.** Loss of *CASP8* enhances radiation killing by Birinapant or Birinapant plus Z-VAD-FMK through induction of necroptosis in HNSCC.

**Figure S8.** Necroptosis sensitivity in HNSCC cell lines

**Figure S9.** Validation of *in vitro* and *in vivo CASP8* knockdown using Tetracycline-Regulated Inducible RNA interference (RNAi) system

**Figure S10.** Treatment schema for *in vivo* experiments

**Figure S11.** Inhibition of *CASP8* function with Emricasan increases radiation killing by Birinapant in HNSCCs.

**Figure S12.** Inhibition of *CASP8* function with Emricasan enhances radiosensitizing effects of Birinapant in HNSCCs.

Supplementary Figure S1.

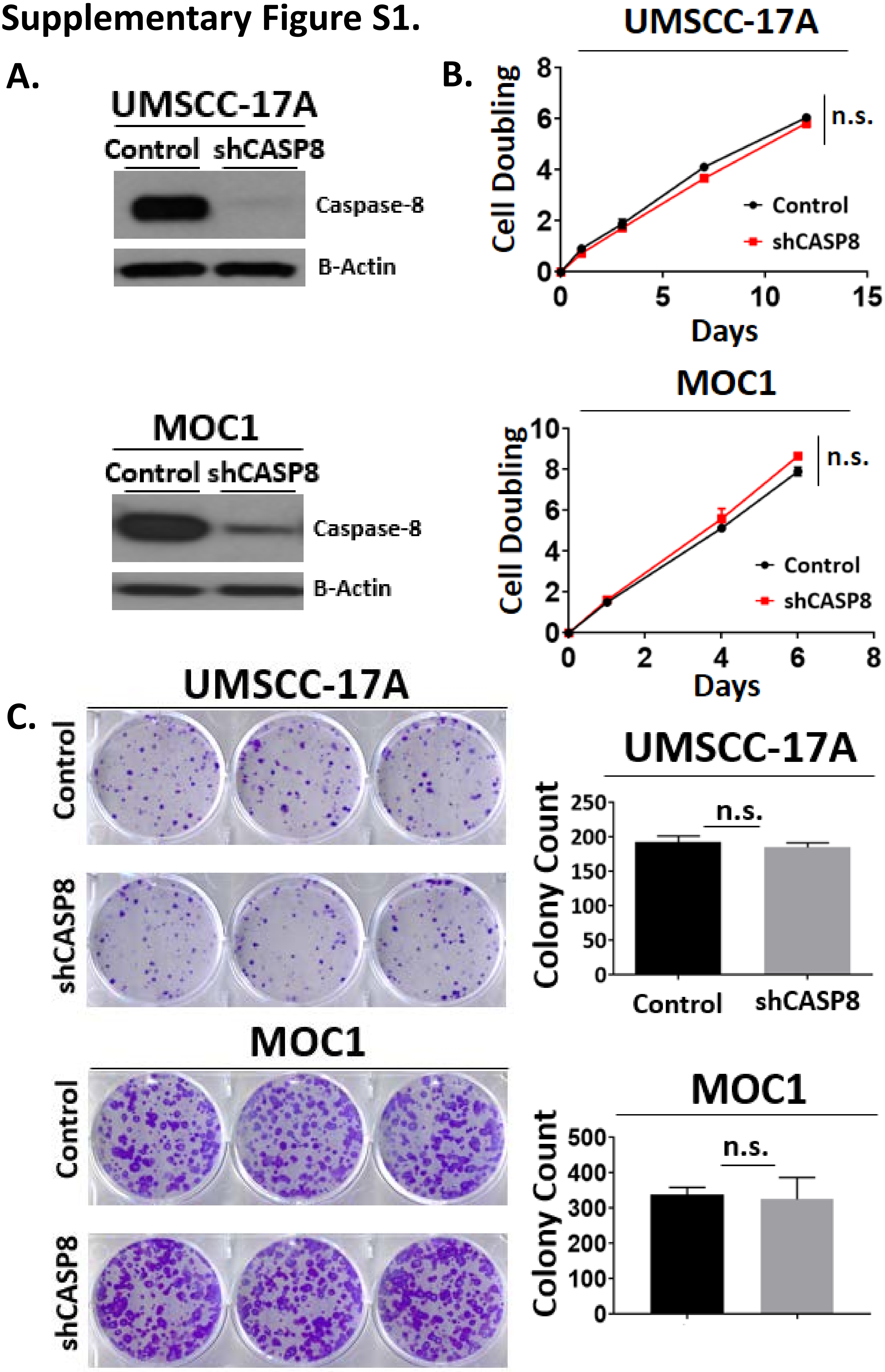

**Supplementary Figure S1. Effects of knockdown of *CASP8* on *in vitro* cell proliferation and clonogenicity in HNSCCs**

**A.** *CASP8* was knockdown using shRNA in UMSCC-17A and MOC1 HNSCC cell lines. Cell lysates obtained from the engineered control (scrambled shRNA) and shCASP8 cell lines were subjected to WB analysis for the validation of *CASP8* knockdown. β-Actin was used as loading control. **B.** Control and shCASP8 UMSCC-17A and MOC1 cell lines were subjected to cell proliferation analysis by CellTiter-Glo. Luminescence reads were taken at the indicated time points and normalized to Day 0 reads to calculate cell doubling. Samples were run in triplicates. Student *t* test was used for statistics. **C.** Control and shCASP8 UMSCC-17A and MOC1 cell lines were subjected to clonogenic survival analysis. Colony counts were used to assess baseline clonogenicity. Samples were run in triplicates. Student *t* test was used for statistics.

### Supplementary Figure S2.

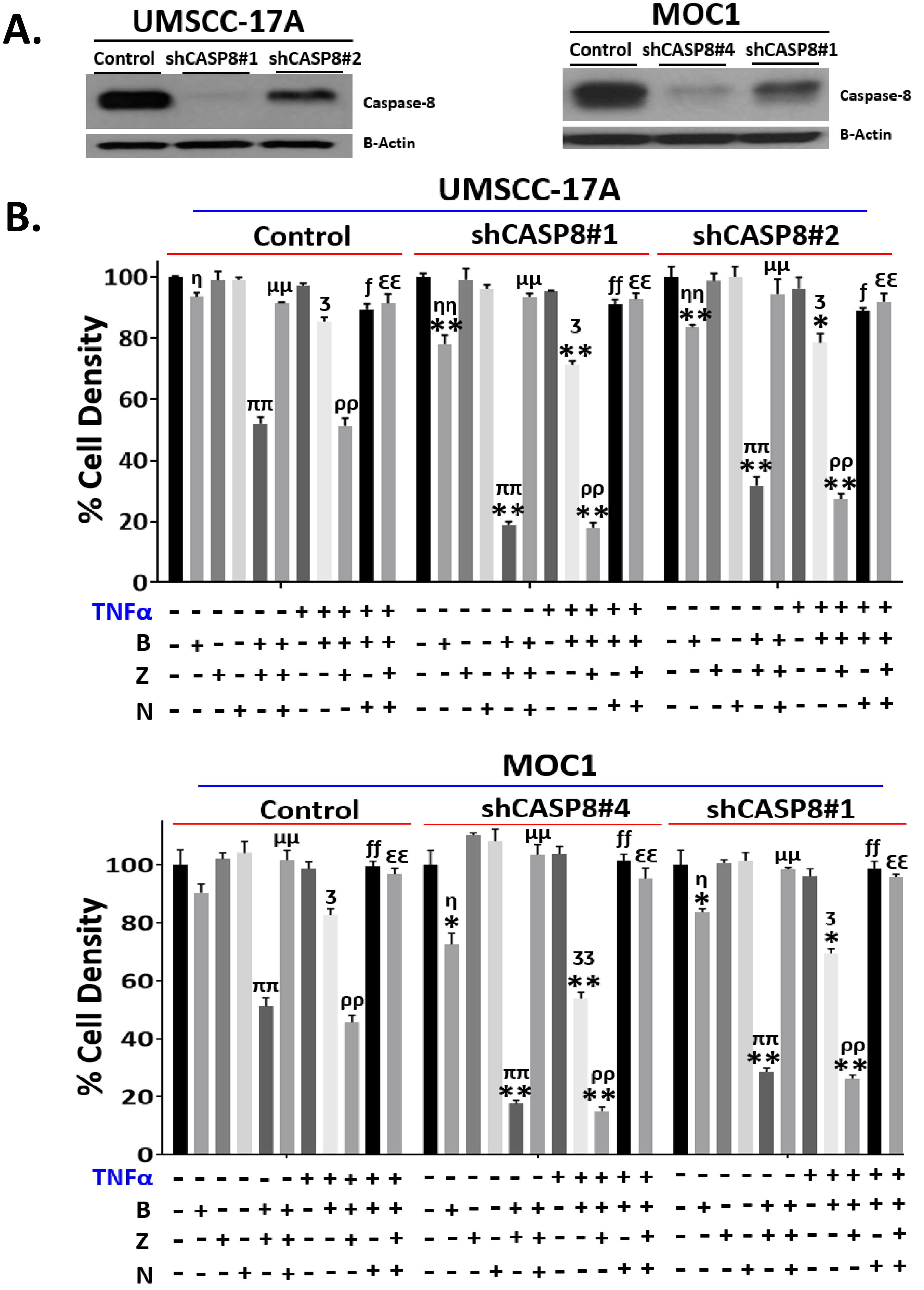


*Legend on following page*

**Supplementary Figure S2. Effects of treatment with Birinapant and Z-VAD-FMK with or without TNFα on cell viability in HNSCCs under loss of *CASP8***

**A.** *CASP8* was knockdown using shRNA in UMSCC-17A and MOC1 HNSCC lines using 2 independent shRNA constructs. Knockdown of *CASP8* was validated by WB analysis. **B.** UMSCC-17A and MOC1 control and shRNA knockdown (shCASP8) cells were treated with Birinapant (**B** [200nmol/L for the UMSCC-17A cells; 1μmol/L for the MOC1 cells]), Z-VAD-FMK (**Z** [5μmol/L for both the cell lines]), Necrostatin-1s (**N** [10μmol/L for both the cell lines]) in the presence and absence of TNFα (**T** [50ng/mL for both the cell lines]) or the combinations as indicated for 24 hours. Cell viability was assessed using CellTiter-Glo. Values normalized to nontreated cells from the same experiment to calculate % cell density (This supplementary figure is related to **Fig. 2A).** All treatments were carried out in triplicates. Student *t* test was used for statistics. *****, *P*<0.05; ******, *P*<0.001 when comparing the effects of B, B+Z , T+B or T+B+Z conditions between isogenic control and shCASP8 cell lines. <u>The following symbols are used to make comparisons between the indicated treatment conditions for each individual cell line</u>: **η**, *P*<0.05; **ηη**, *P*<0.001 to compare no treatment (NT) vs B. **Ʒ,** *P*<0.05; **ƷƷ,** *P*<0.001 to compare B vs T+B. ***f***, *P*<0.05; ***ff**, P*<0.001 to compare T+B vs T+B+N. **Ɛ,** P<0.05; **ƐƐ,** *P*<0.001 to compare T+B+Z vs T+B+Z+N. **μ,** *P*<0.05; **μμ,** *P*<0.001 to compare B+Z vs B+Z+N. **π,** *P*<0.05; **ππ,** *P*<0.001 to compare B vs B+Z. **ρ,** *P*<0.05; **ρρ,** *P*<0.001 to compare T+B vs T+B+Z.

### Supplementary Figure S3.

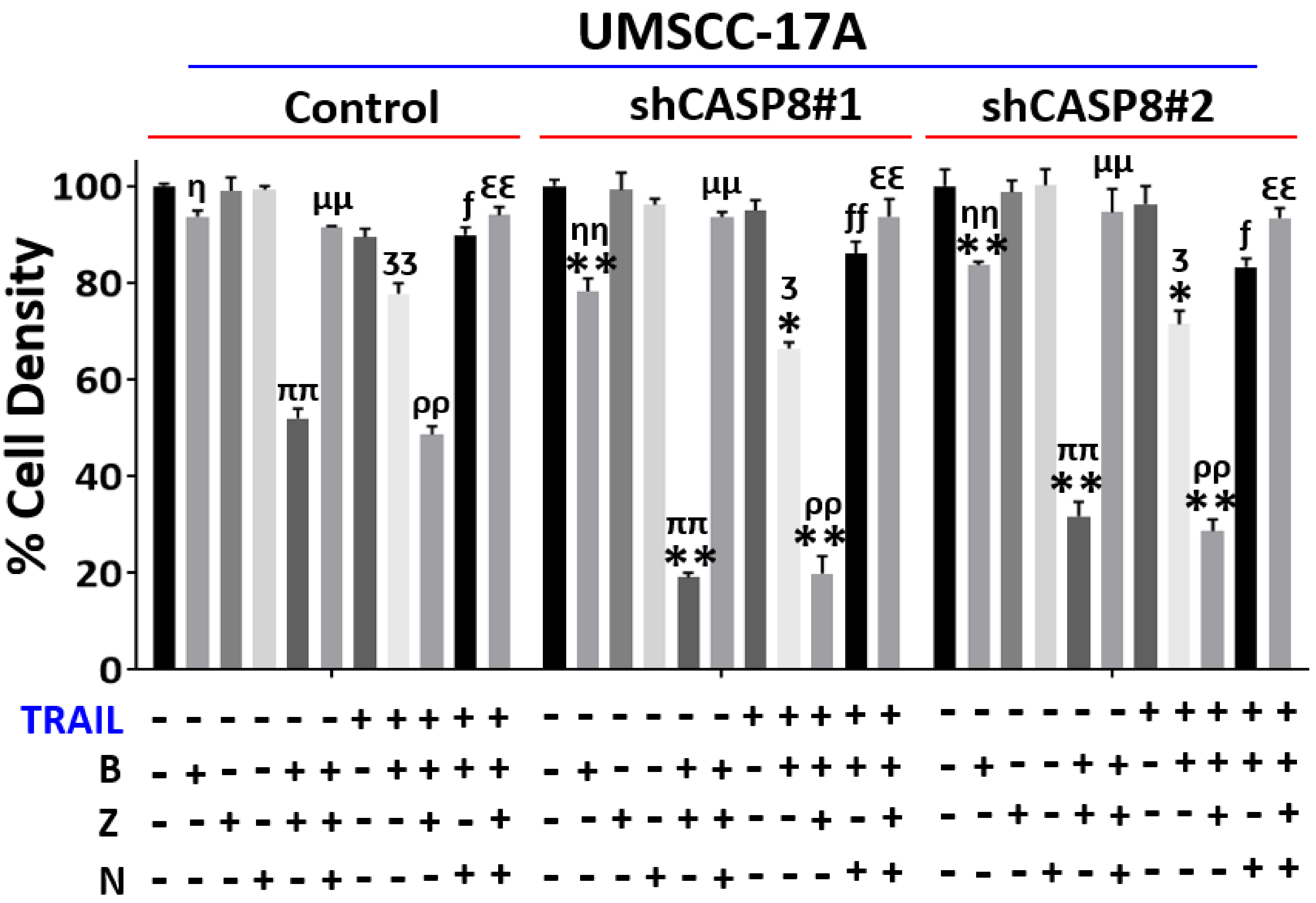


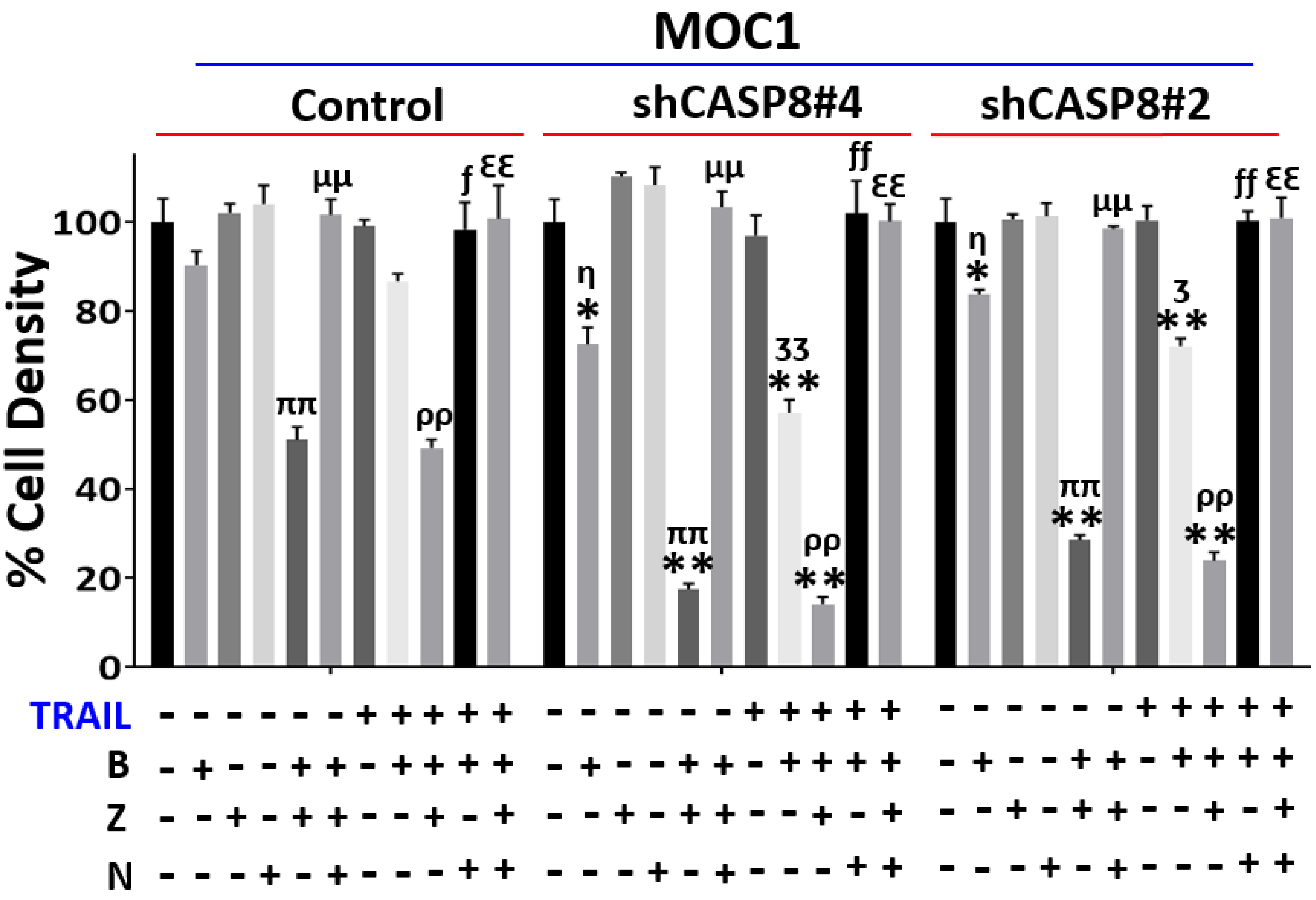


*Legend on following page*

**Supplementary Figure S3. Effects of treatment with Birinapant and Z-VAD-FMK with or without TRAIL on cell viability in HNSCCs under loss of *CASP8***

**A.** *CASP8* was knockdown using shRNA in UMSCC-17A and MOC1 HNSCC lines using 2 independent shRNA constructs. Knockdown of *CASP8* was validated by WB analysis. **B.** UMSCC-17A and MOC1 control and shRNA knockdown (shCASP8) cells were treated with Birinapant (**B** [200nmol/L for the UMSCC-17A cells; 1μmol/L for the MOC1 cells]), Z-VAD-FMK (**Z** [5μmol/L for both the cell lines]), Necrostatin-1s (**N** [10μmol/L for both the cell lines]) in the presence and absence of TRAIL (**T** [10ng/ml for the UMSCC-17A cells; 50ng/mL for the MOC1 cells]) or the combinations as indicated for 24 hours. Cell viability was assessed using CellTiter-Glo. Values normalized to nontreated cells from the same experiment to calculate % cell density (This supplementary figure is related to **Fig. 2A).** All treatments were carried out in triplicates. Student *t* test was used for statistics. *****, $P<0.05$; ******, $P<0.001$ when comparing the effects of B, B+Z , T+B or T+B+Z conditions between isogenic control and shCASP8 cell lines. <u>The following symbols are used to make comparisons between the indicated treatment conditions for each individual cell line</u>: **η**, $P<0.05$; **ηη**, $P<0.001$ to compare no treatment (NT) vs B. **ʒ,** $P<0.05$; **ʒʒ**,$P<0.001$ to compare B vs T+B. ***ƒ***, $P<0.05$; ***ƒƒ**,* $P<0.001$ to compare T+B vs T+B+N. **ɛ,** P<0.05; **ɛɛ,** $P<0.001$ to compare T+B+Z vs T+B+Z+N. **μ,** $P<0.05$; **μμ,** $P<0.001$ to compare B+Z vs B+Z+N. **π,** $P<0.05$; **ππ,** $P<0.001$ to compare B vs B+Z. **ρ,** $P<0.05$; **ρρ,** $P<0.001$ to compare T+B vs T+B+Z.

#### Supplementary Figure S4.

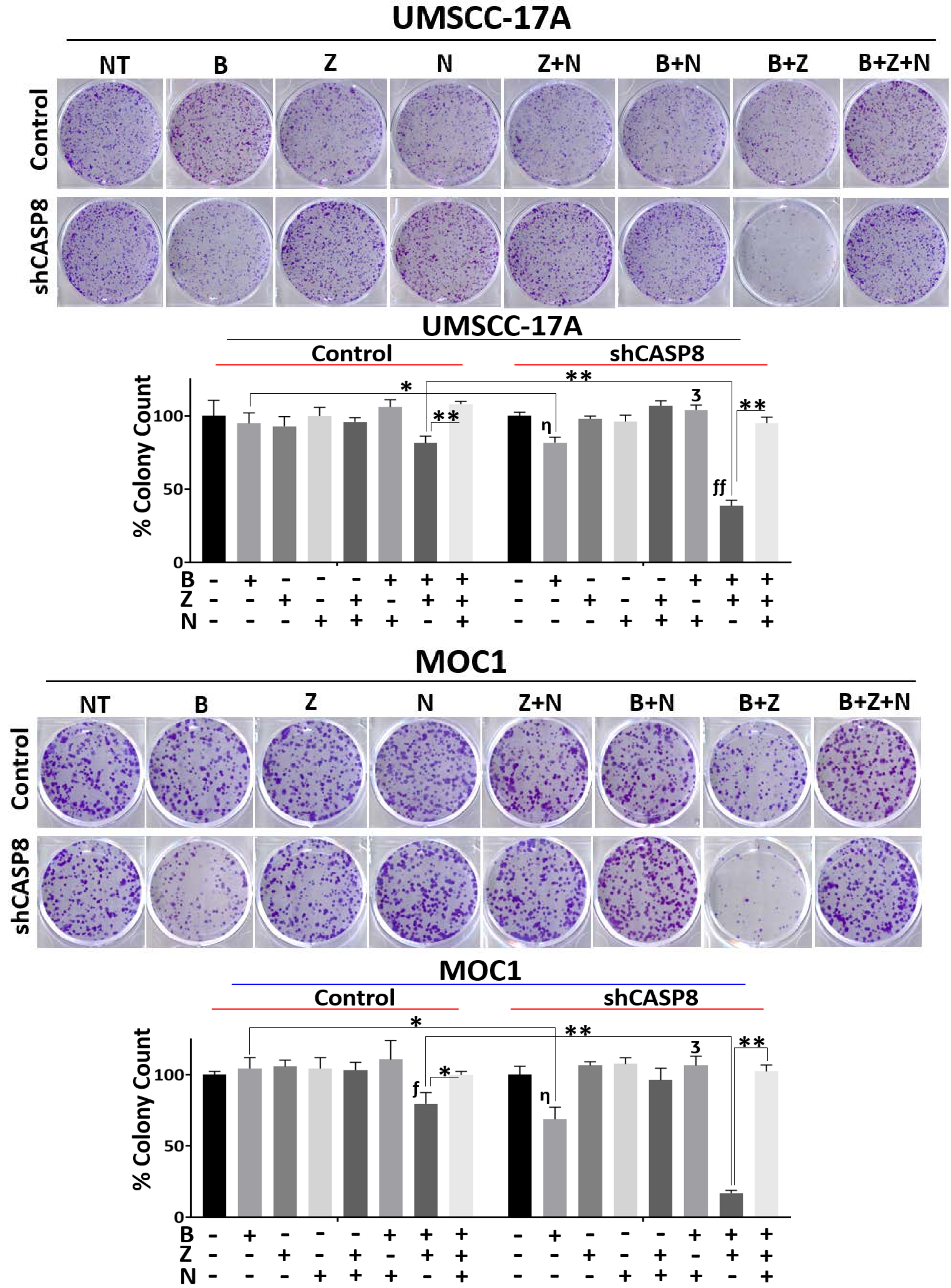


*Legend on following page*

**Supplementary Figure S4. Effects of treatment with Birinapant and Z-VAD-FMK on clonogenicity in HNSCCs under loss of *CASP8***

UMSCC-17A and MOC1 control and shRNA knockdown (shCASP8) cells were treated with Birinapant (**B** [50nmol/L for the UMSCC-17A cells; 250nmol/L for the MOC1 cells]), Z-VAD-FMK (**Z** [5μmol/L for both the cell lines]), Necrostatin-1s (**N** [10μmol/L for both the cell lines]) or the combinations as indicated. 24 hour after treatments, drug dilutions were washed out, colonies were allowed to form for 5-12 days, after which they were stained and counted. Surviving colonies were normalized to nontreated cells from the same experiment to calculate % colony count (This supplementary figure is related to **Fig. 2B).** All treatments were carried out in triplicates. Student *t* test was used for statistics. *****, *P*<0.05; ******, *P*<0.001 for the indicated pairwise comparisons. The following symbols are used to make comparisons between the indicated treatment conditions for each individual cell line: **η**, *P*<0.05; **ηη**, *P*<0.001 to compare no treatment (NT) vs B. **ʒ,** *P*<0.05; **ʒʒ**, *P*<0.001 to compare B vs B+N. ***f***, *P*<0.05; ***ff***, *P*<0.001 to compare B vs B+Z.

### Supplementary Figure S5.

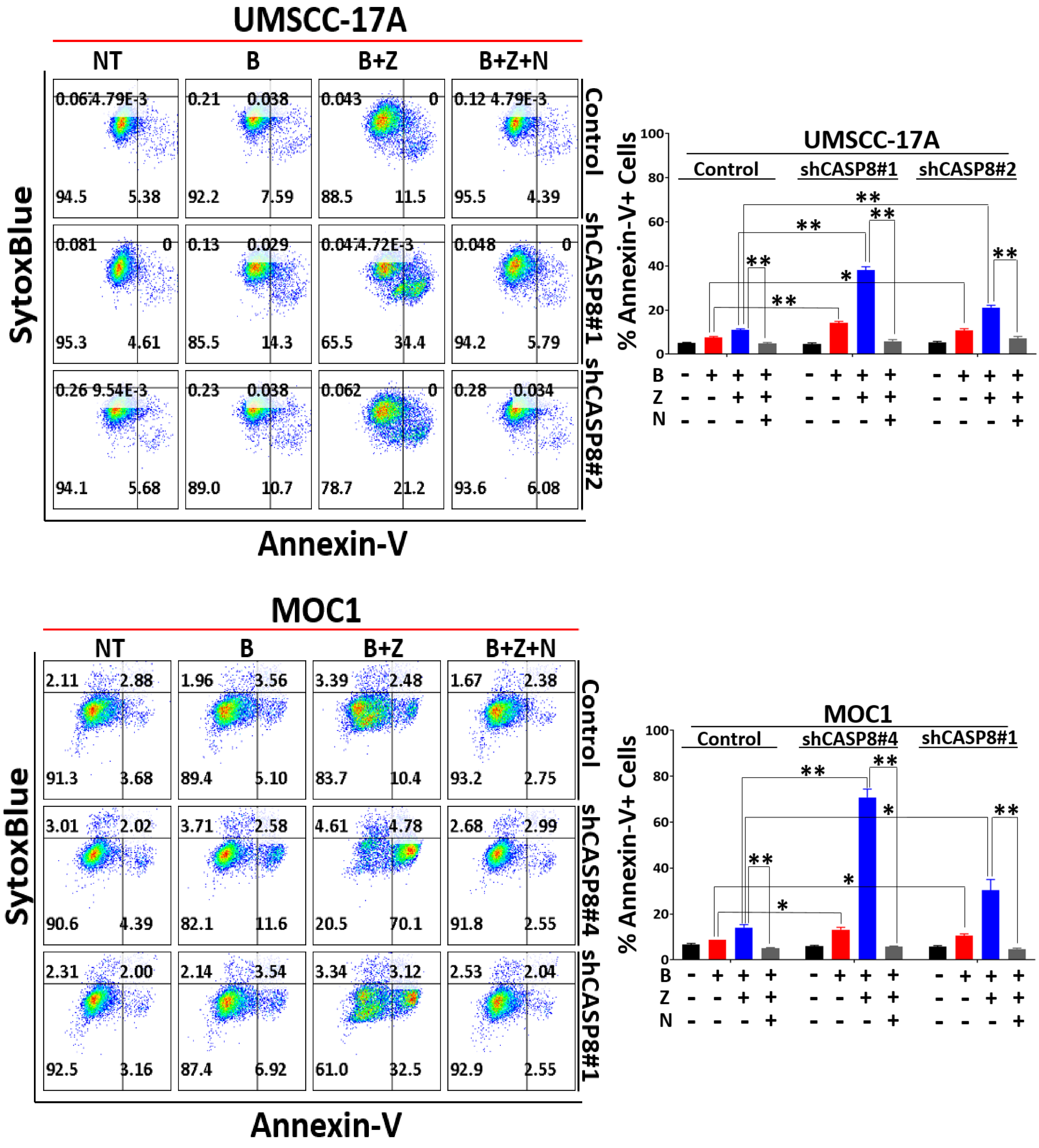


*Legend on following page*

**Supplementary Figure S5. Necroptotic effects of treatment with Birinapant and Z-VAD-FMK are enhanced under loss of *CASP8* in HNSCCs.**

UMSCC-17A and MOC1 control and shRNA knockdown (shCASP8) cells (2 independent shRNA clones for each cell line) were treated with Birinapant (**B** [200nmol/L for the UMSCC-17A cells; 1μmol/L for the MOC1 cells]), Z-VAD-FMK (**Z** [5μmol/L for both the cell lines]), Necrostatin-1s (**N** [10μmol/L for both the cell lines]) or the combinations as indicated. AnnexinV-APC/SytoxBlue staining was performed 24 hour after treatments. % Annexin-V positivity was used as a measure to assess cell death (This supplementary figure is related to **Fig. 2C**). All treatments were carried out in triplicates. Student *t* test was used for statistics. *, $P<0.05$; **, $P<0.001$ for the indicated pairwise comparisons.

#### Supplementary Figure S6.

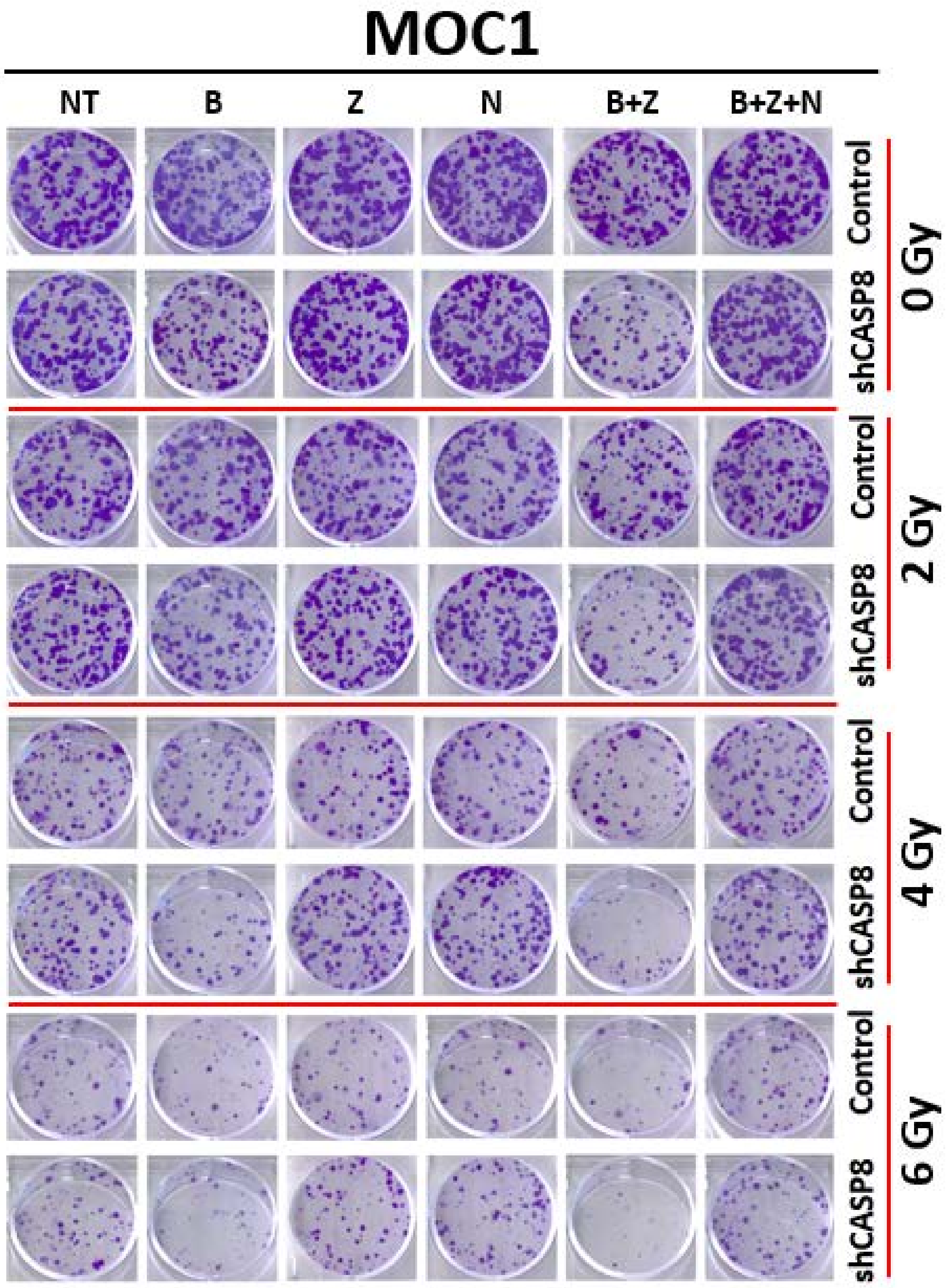


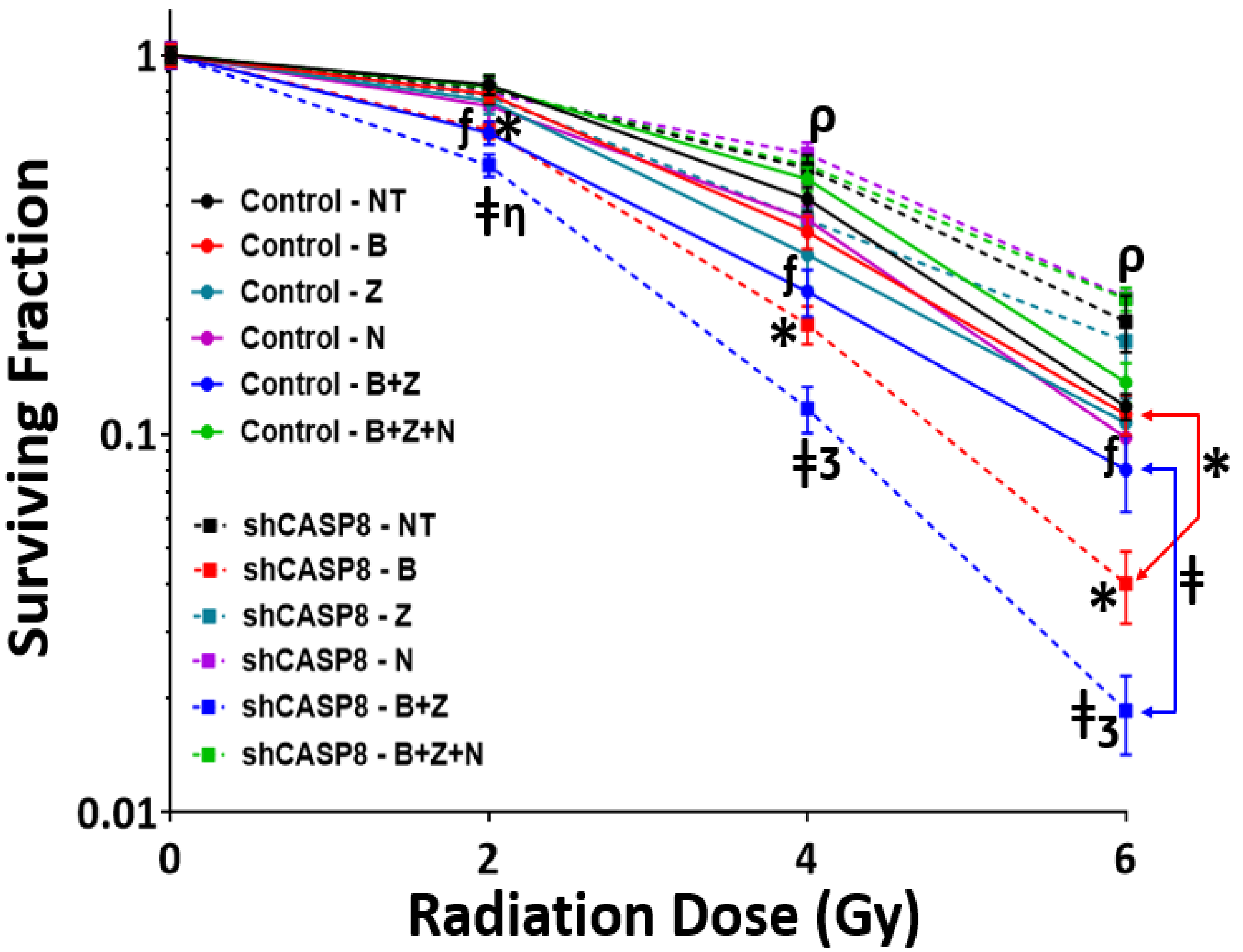


*Legend on following page*

**Supplementary Figure S6. Loss of *CASP8* increases the radiosensitizing effects of Birinapant or Birinapant plus Z-VAD-FMK through induction of necroptosis in HNSCC.**

MOC1 control and shCASP8 cells were treated with radiation (**X** [2, 4 and 6 Gy]), Birinapant (**B** [125nmol/L]), Z-VAD-FMK (**Z** [5μmol/L]), Necrostatin-1s (**N** [10μmol/L]) or the combinations as indicated. 24 hour after treatments, drug dilutions were washed out, colonies were allowed to form for 5 days, after which they were stained and counted. Surviving colony counts were normalized to nontreated cells (cells treated with no drugs) of each radiation dose from the same experiment. *Log10* of surviving fractions were plotted (This supplementary figure is related to **Fig. 3A**). All treatments were carried out in triplicates. Student *t* test was used for statistics. *, *P*<0.05; when comparing surviving fractions following X+B treatments between control and shCASP8 cells **ǂ,** *P*<0.05; when comparing surviving fractions following X+B+Z treatments between control and shCASP8 cells ***ƒ*****,** *P*<0.05; when showing reversal of death upon addition of N to X+B+Z for the control cells, **η,** *P*<0.05; when showing reversal of death upon addition of N to X+B+Z for the shCASP8 cells, **ʒ,** *P*<0.001; when showing reversal of death upon addition of N to X+B+Z for the shCASP8 cells. Symbols are placed at the radiation doses they refer to.

#### Supplementary Figure S7.

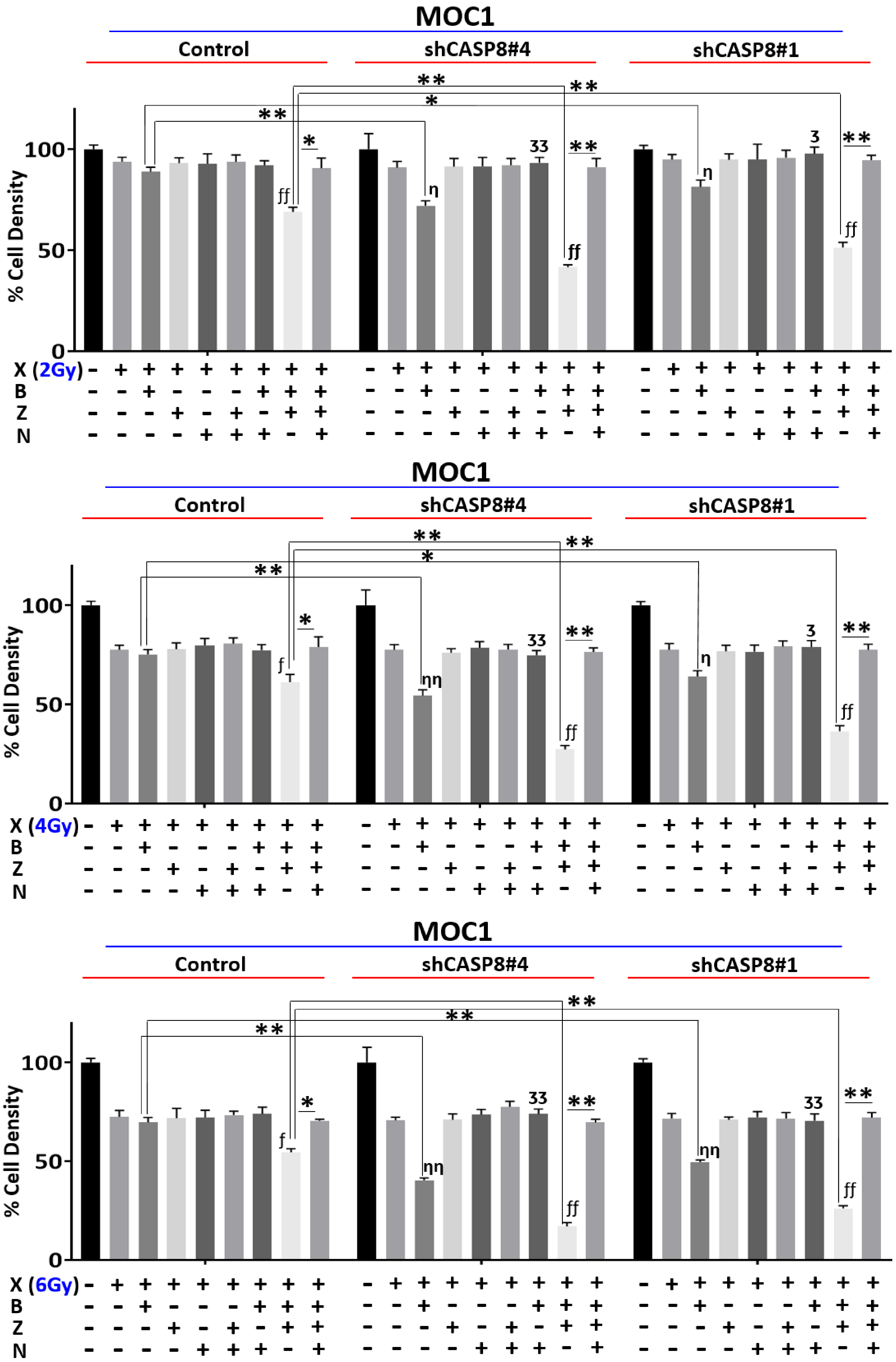


*Legend on following page*

**Supplementary Figure S7. Loss of *CASP8* enhances radiation killing by Birinapant or Birinapant plus Z-VAD-FMK through induction of necroptosis in HNSCC.**

MOC1 control and shCASP8 cells were treated with radiation (**X** [2, 4 and 6 Gy]), Birinapant (**B** [250nmol/L]), Z-VAD-FMK (**Z** [5μmol/L]), Necrostatin-1s (**N** [10μmol/L]) or the combinations as indicated. 24 hour after treatments, cell viability was assessed by Cell-Titer Glo. Values normalized to nontreated cells from the same experiment to calculate % cell density (This supplementary figure is related to **Fig. 3B**). All treatments were carried out in triplicates. Student t test was used for statistics. *, P<0.05; **, P<0.001 for the indicated pairwise comparisons. <u>The following symbols are used to make comparisons between the indicated treatment conditions for each individual cell line</u>: **η**, P<0.05; **ηη,** P<0.001 to compare X vs X+B. **ʒ,** P<0.05; **ʒʒ,** P<0.001 to compare X+B vs X+B+N. ***ƒ*,** P<0.05; ***ƒƒ*,** P<0.001 to compare X vs X+B+Z.

#### Supplementary Figure S8.

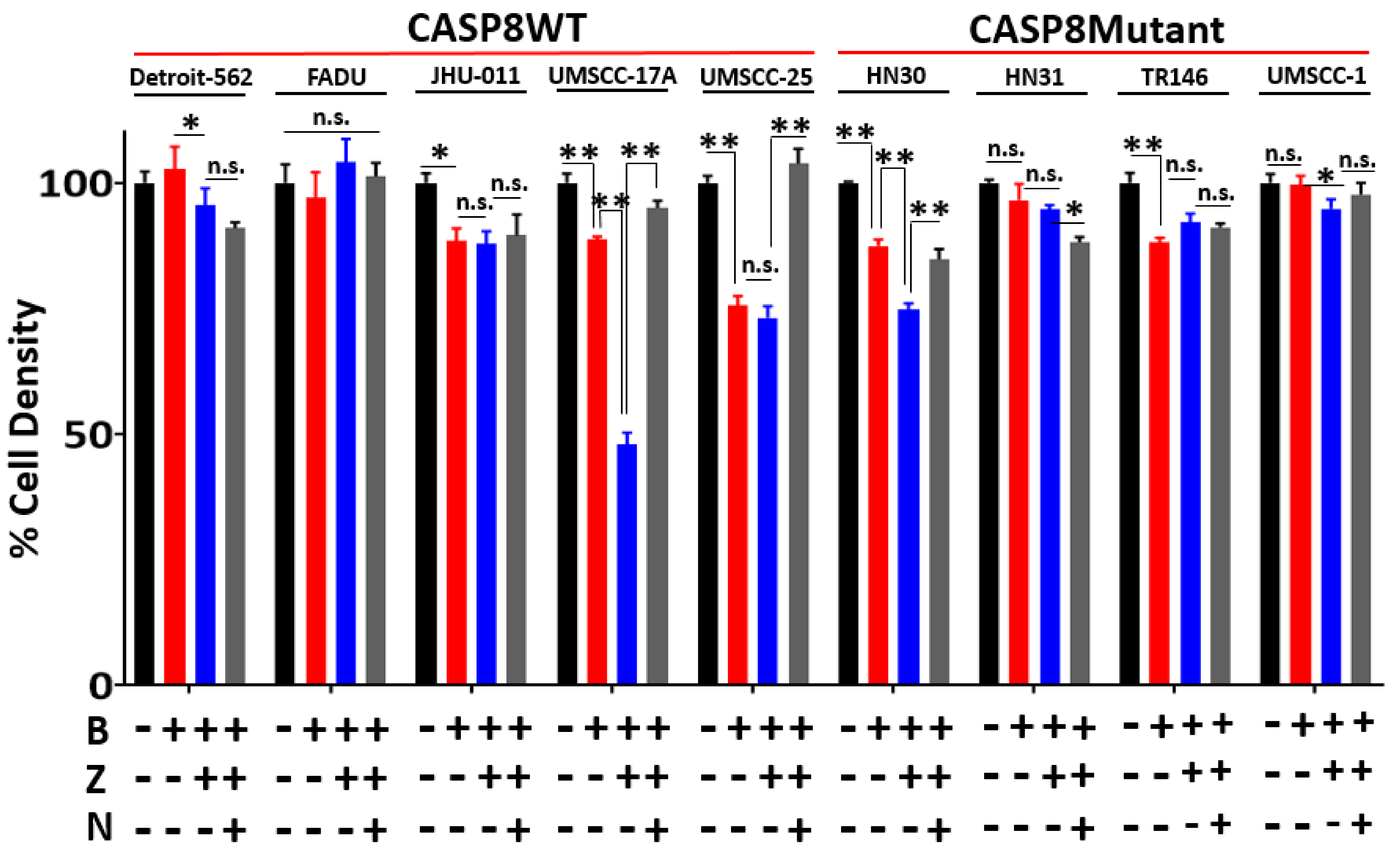


*Legend on following page*

**Supplementary Figure S8. Necroptosis sensitivity in HNSCC cell lines**

**A.** A panel of **9** human-derived HNSCC cell lines, **5** *CASP8* WT (Detroit-562, FADU, JHU-011, UMSCC-17A and UMSCC-25) and **4** *CASP8* mutant (HN30, HN31, TR146 and UMSCC-1) were treated with Birinapant (**B** [1μmol/L]), Z-VAD-FMK (**Z** [5μmol/L]), Necrostatin-1s (**N** [10μmol/L]) or the combinations. 24 hour after treatments, cell viability was assessed by Cell-Titer Glo. Values normalized to nontreated cells from the same experiment to calculate % cell density (This supplementary figure is related to **Fig. 5A**). All treatments were carried out in triplicates. Student t test was used for statistics. *, $P<0.05$; **, $P<0.001$ for the indicated pairwise comparisons.

### Supplementary Figure S9.

A.

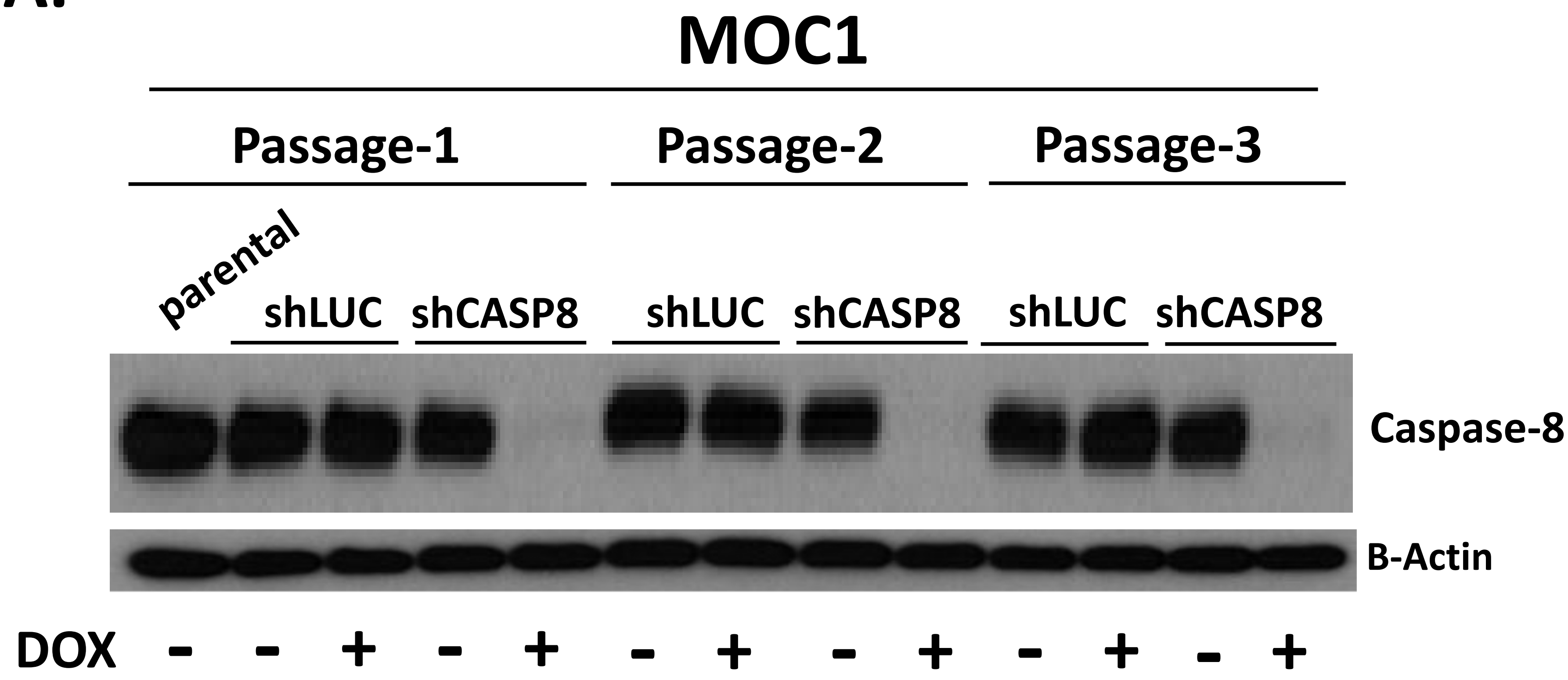


B.

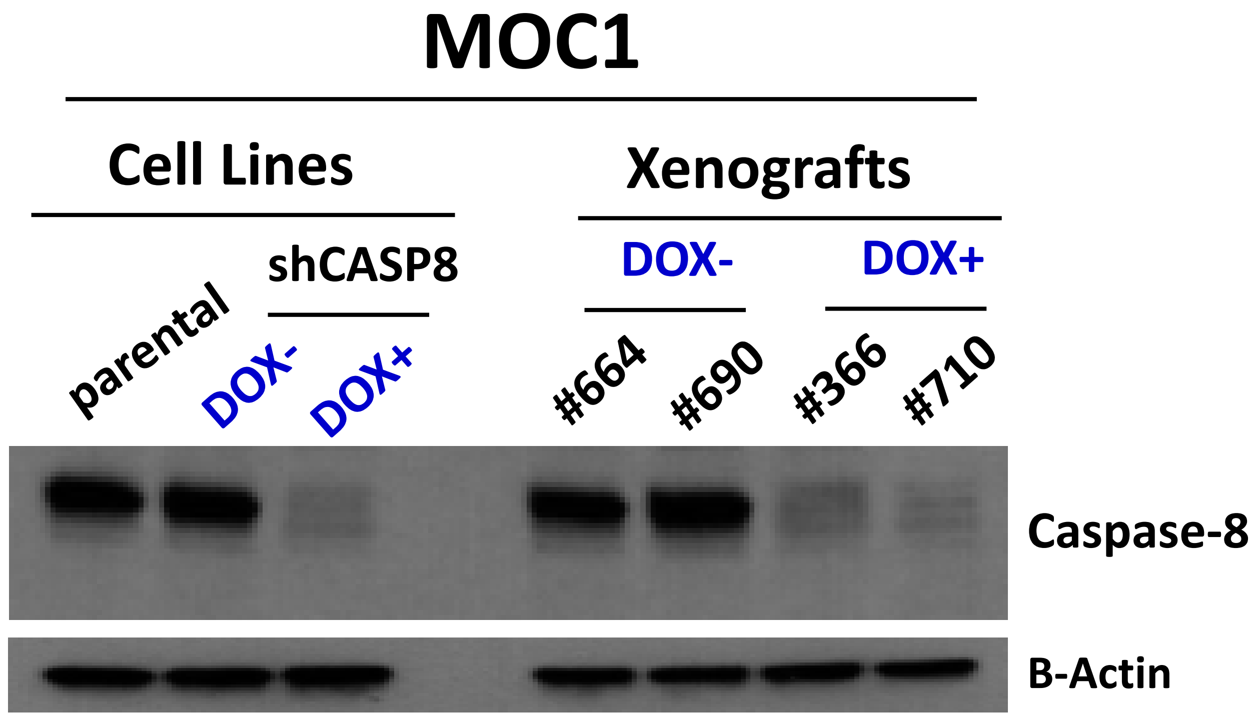


**Supplementary Figure S9. Validation of *in vitro* and *in vivo* *CASP8* knockdown using Tetracycline-Regulated Inducible RNA interference (RNAi) system**

This supplementary figure is related to **Fig 6A-B.** **A.** MOC1 cells were transduced with lentiviral constructs designed against Luciferase (shLUC) or *CASP8 (*shCASP8*)* (33). The engineered shLUC and shCASP8 cells were cultured in the presence and absence of Doxycycline (50ng/ml). 72 hours after treatment, cells were passaged and cell lysates were obtained. This cycle was repeated 3 times. Cell lysates obtained from each cycle along with that from MOC1 parental cells were subjected to WB analysis for *CASP8*. β-Actin was used as loading control. **B.** Tumor samples collected from control (#664, #690) and Doxycycline (DOX)-fed (#366, #710) mice were minced and cultured in medium for 48 hours. Cells shed from the indicated xenografts that have attached to culture dishes were collected and lysed. Cell lysates obtained from the tumor samples along with those from the indicated cell lines were subjected to WB analysis for *CASP8*. β-Actin was used as loading control.

#### Supplementary Figure S10.

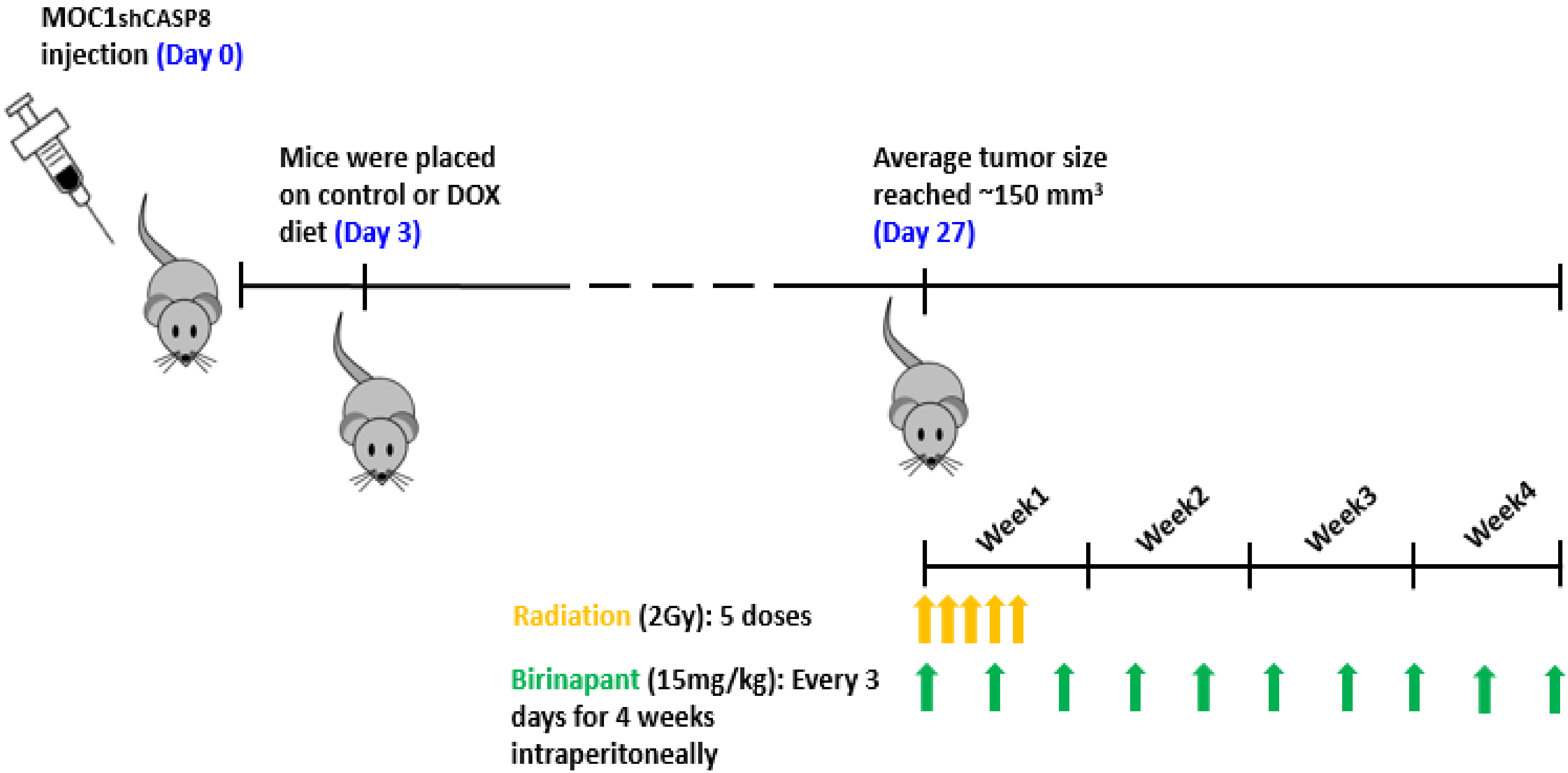


*Legend on following page*

**Supplementary Figure S10. Treatment schema for *in vivo* experiments**

This supplementary figure is related to **Fig 6A-B.**

**Procedure shown on the treatment schema:** $2 \times 10^6$ MOC1 cells transduced with an inducible shRNA against *CASP8* were injected into the right flank of WT female C57BL/6 mice. Mice were randomized and placed on control or DOX diet (doxycycline hyclate added at 625 mg/kg) 3 days post injection to induce knockdown of *CASP8 in vivo* (Please refer to **Supplementary Fig. S9** for the WB images). Control and *CASP8* knockdown mice were randomized into 4 treatment groups (vehicle control, 15mg/kg Birinapant, 5X2Gy radiation or combination, *n*=7-10/each) 27 days post inoculation when the average tumor volume reached ~150 $mm^3$. Radiation started on Day 27: 2Gy of radiation given Monday to Friday for 1 week. Birinapant started on Day 27: 15mg/kg Birinapant given intraperitoneally every 3 days for 4 weeks.

#### Supplementary Figure S11.

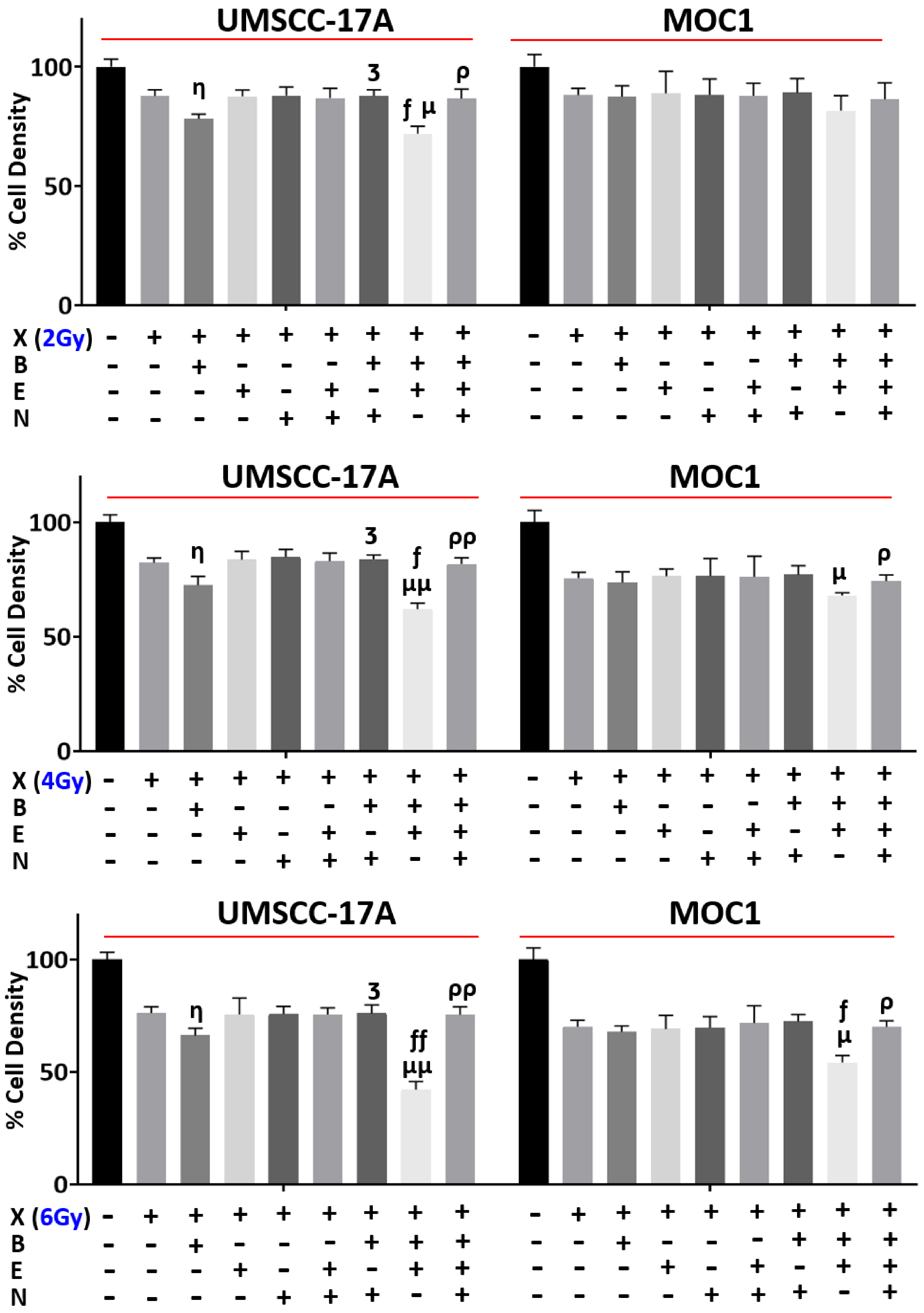


*Legend on following page*

**Supplementary Figure S11. Inhibition of *CASP8* function with Emricasan increases radiation killing by Birinapant in HNSCCs.**

MOC1 and UMSCC-17A parental cells were treated with radiation (**X** [2, 4 and 6 Gy for both the cell lines]), Birinapant (**B** [50nmol/L for the UMSCC-17A cells and 250nmol/L for the MOC1 cells]), Emricasan (**E** [1μmol/L for both the cell lines]), Necrostatin-1s (**N** [10μmol/L for both the cell lines]) or the combinations as indicated. 24 hour after treatments, cell viability was assessed by Cell-Titer Glo. Values normalized to nontreated cells from the same experiment to calculate % cell density (This supplementary figure is related to **Fig. 6C**). All treatments were carried out in triplicates. Student t test was used for statistics. The following symbols are used to make comparisons between the indicated treatment conditions for each individual cell line: **η**, P<0.05; **ηη,** P<0.001 to compare X vs X+B. **ʒ,** P<0.05; **ʒʒ,** P<0.001 to compare X+B vs X+B+N. ***ƒ*,** P<0.05; ***ƒƒ*,** P<0.001 to compare X+B vs X+B+E. **μ,** P<0.05; **μμ,** P<0.001 X vs X+B+E. **ρ,** P<0.05; **ρρ,** P<0.001 to compare X+B+E vs X+B+E+N.

#### Supplementary Figure S12.

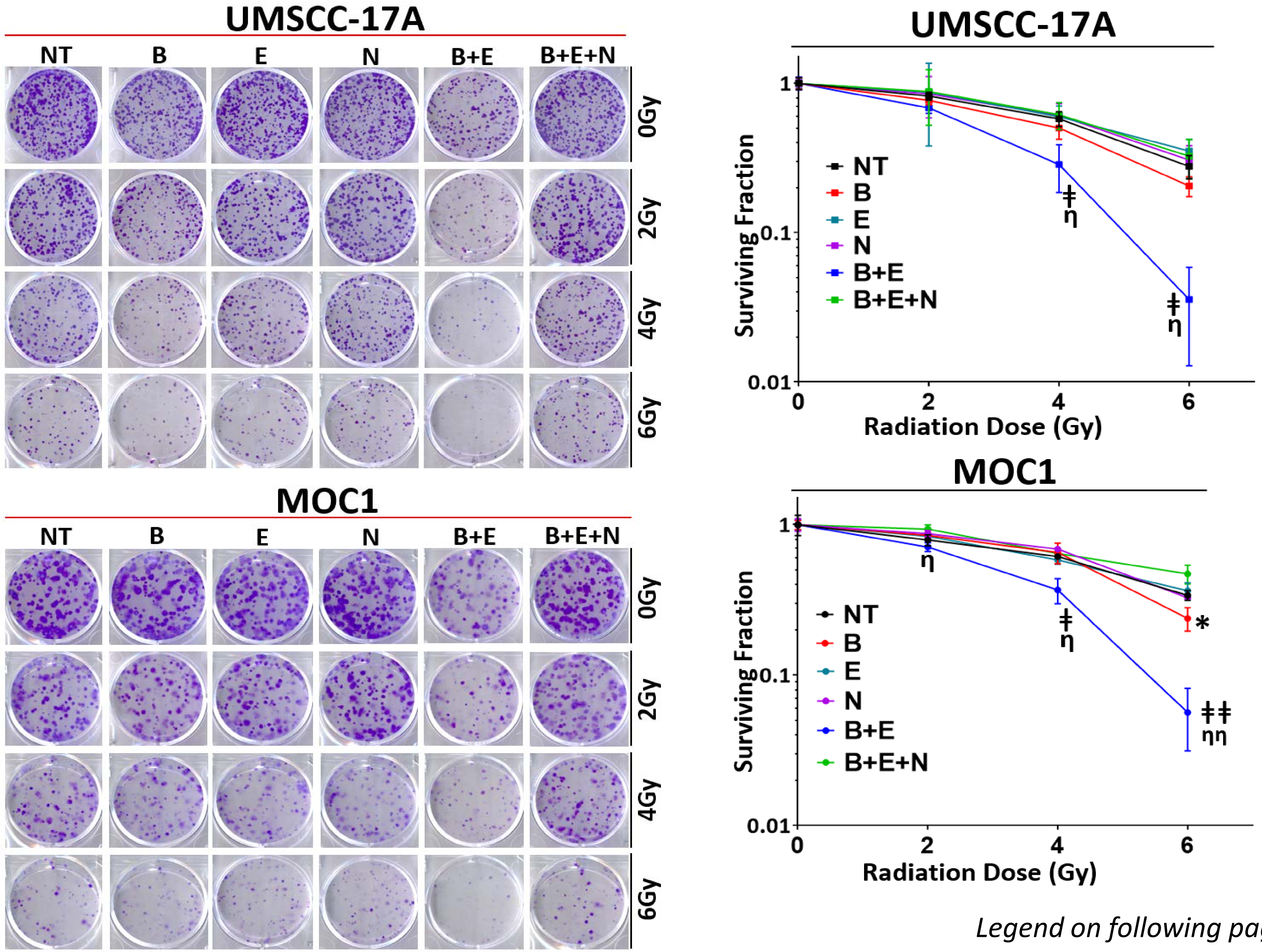


*Legend on following page*

**Supplementary Figure S12. Inhibition of *CASP8* function with Emricasan enhances radiosensitizing effects of Birinapant in HNSCCs.**

MOC1 and UMSCC-17A parental cells were treated with radiation (**X** [2, 4 and 6 Gy for both the cell lines]), Birinapant (**B** [25nmol/L for the UMSCC-17A cells and 125nmol/L for the MOC1 cells]), Emricasan (**E** [1μmol/L for both the cell lines]), Necrostatin-1s (**N** [10μmol/L for both the cell lines]) or the combinations as indicated. 24 hour after treatments, drug dilutions were washed out, colonies were allowed to form for 5-12 days, after which they were stained and counted. Surviving colony counts were normalized to nontreated cells (cells treated with no drugs) of each radiation dose from the same experiment. Log10 of surviving fractions were plotted (This supplementary figure is related to **Fig. 6D**). All treatments were carried out in triplicates. Student *t* test was used for statistics. *, $P<0.05$; when comparing X+B to X alone for the indicated radiation dose. **ǂ,** $P<0.05$ and **ǂǂ,** $P<0.001$; when comparing X+B+E to X alone for the indicated radiation doses. **η,** $P<0.05$ and **ηη,** $P<0.001$; when comparing X+B+E to X+B+E+N for the indicated radiation doses.
