## Supplemental tables for "Loss of Caspase-8 function in combination with SMAC mimetic treatment sensitizes Head and Neck Squamous Carcinoma to radiation through induction of necroptosis"

**Supplementary Table S1: shRNA and CRIPSR-Cas9 sgRNA oligo sequences**

| mouse *Casp8* CRISPR-Cas9 sgRNA sequences | DNA Sequence (5' to 3') |
|---|---|
| *Casp8* for-1 | CACCGGTGTCGTCTATGGAACGGAT |
| *Casp8* rev-1 | AAACATCCGTTCCATAGACGACACC |
| *Casp8* for-2 | CACCGCAAGAACTATATTCCGGATG |
| *Casp8* rev-2 | AAACCATCCGGAATATAGTTCTTGC |
| *Casp8* for-3 | CACCGTAGCTTCTGGGCATCCTCGA |
| *Casp8* rev-3 | AAACTCGAGGATGCCCAGAAGCTAC |
| **human *CASP8* shRNA oligo sequences** | |
| *CASP8* shRNA#1 (shCASP8#1) | TAATTCGGAAGAGCAGCTC |
| *CASP8* shRNA#1 (shCASP8#2) | TTCCTTCTCCCAGGATGAC |
| **mouse *Casp8* shRNA oligo sequences** | |
| *Casp8* shRNA#1 (shCASP8#1) | AAACTTTGTCTGAAGTCCG |
| *Casp8* shRNA#4 (shCASP8#4) | TTTCATTTGCAGTGCAGTC |
| **mouse *Casp8* inducible shRNA oligo sequences** | |
| *Casp8* inducible shRNA#10 (shCASP8) | TTTCATTTGCAGTGCAGTC |
| **Luciferase inducible shRNA oligo sequences** | |
| luciferase inducible shRNA (shLUC) | CGCTGAGTACTTCGAAATGTC |
| **mouse *Ripk3* shRNA oligo sequences** | |
| *Ripk3* shRNA (shRIP3) | TACCTCGGAGACAGCAGCA |

### Supplementary Table S2: List of reagents/drugs used in the study

| No. | Reagent/drug name | Catalog number | Company |
|---|---|---|---|
| 1 | Birinapant | S7015 | Selleckchem |
| 2 | zVAD-FMK | FMK001 | R&D Systems |
| 3 | human recombinant TNFα | 210-TA | R&D Systems |
| 4 | human recombinant TRAIL | 375-TEC | R&D Systems |
| 5 | murine recombinant TNFα | 315-01A | Peprotech |
| 6 | murine recombinant TRAIL | 315-19 | Peprotech |
| 7 | Necrostatin-1s (7-Cl-O-Nec1) | 10-4544 | Focus Biomolecules |
| 8 | Puromycin dihydrochloride from Streptomyces alboniger | P8833 | Sigma-Aldrich |
| 9 | Doxycycline | 631311 | Takara Bio USA |

### Supplementary Table S3: List of antibodies used for western blotting

| No. | Antibody name | Catalog number | Source | Company | Dilution |
|---|---|---|---|---|---|
| 1 | human CASP8 | 551242 | mouse | BD Biosciences | 1:1000 |
| 2 | human/mouse CASP8 | AF1650 | rabbit | R&D Systems | 1:400 |
| 3 | RIP1 | 610458 | mouse | BD Biosciences | 1:1000 |
| 4 | human RIP3 (E1Z1D) | 13526 | rabbit | Cell Signaling | 1:1000 |
| 5 | mouse RIP3 (D4G2A) | 95702 | rabbit | Cell Signaling | 1:1000 |
| 6 | MLKL (D216N) | 14993 | rabbit | Cell Signaling | 1:1000 |
| 7 | MLKL (mouse specific) | 28640 | rabbit | Cell Signaling | 1:1000 |
| 8 | phospho-RIP1 (S166) | 31122 | rabbit | Cell Signaling | 1:1000 |
| 9 | phospho-MLKL (S358) (D6H3V) | 91689 | rabbit | Cell Signaling | 1:1000 |
| 10 | PARP | 9542 | rabbit | Cell Signaling | 1:1000 |
| 11 | Anti-HA | H3663 | mouse | Sigma-Aldrich | 1:2000 |
| 12 | β-actin | A1978 | mouse | Sigma-Aldrich | 1:10000 |
| 13 | Anti-rabbit IgG | 1706515 | goat | BioRad | 1:5000 |
| 14 | Anti-mouse IgG | 1706516 | goat | BioRad | 1:5000 |

**Supplementary Table S4: *p* values for key pairwise comparisons in <u>tumor growth analysis</u>**

| Compared Animal Cohorts | | *p* value |
|---|---|---|
| Cohort#1 | Cohort#2 | |
| Control-NT | Control-B | 0.7628 |
| Control-NT | Control-X | 0.0093 |
| Control-NT | Control-X+B | <0.001 |
| Control-B | Control-X | 0.0795 |
| Control-B | Control-X+B | <0.001 |
| Control-X | Control-X+B | 0.1357 |
| shCASP8-NT | shCASP8-B | <0.001 |
| shCASP8-NT | shCASP8-X | <0.001 |
| shCASP8-NT | shCASP8-X+B | <0.001 |
| shCASP8-B | shCASP8-X | <0.001 |
| shCASP8-B | shCASP8-X+B | <0.001 |
| shCASP8-X | shCASP8-X+B | <0.001 |
| Control-NT | shCASP8-NT | <0.001 |
| Control-B | shCASP8-B | <0.001 |
| Control-X | shCASP8-X | 0.9714 |
| Control-X+B | shCASP8-X+B | <0.001 |

**Supplementary Table S5: *p* values for key pairwise comparisons in <u>survival analysis</u>**

| Compared Animal Cohorts | | *p* value |
|---|---|---|
| Cohort#1 | Cohort#2 | |
| Control-NT | Control-B | 0.4345 |
| Control-NT | Control-X | <0.001 |
| Control-NT | Control-X+B | <0.001 |
| Control-B | Control-X | <0.001 |
| Control-B | Control-X+B | <0.001 |
| Control-X | Control-X+B | 0.0206 |
| shCASP8-NT | shCASP8-B | <0.001 |
| shCASP8-NT | shCASP8-X | <0.001 |
| shCASP8-NT | shCASP8-X+B | <0.001 |
| shCASP8-B | shCASP8-X | <0.001 |
| shCASP8-B | shCASP8-X+B | <0.001 |
| shCASP8-X | shCASP8-X+B | <0.001 |
| Control-NT | shCASP8-NT | 0.4846 |
| Control-B | shCASP8-B | <0.001 |
| Control-X | shCASP8-X | 0.4747 |
| Control-X+B | shCASP8-X+B | <0.001 |
